# Synaptic Input Triggers On-Demand Spine-Specific Mitochondrial ATP Production and Delivery

**DOI:** 10.64898/2026.03.09.708229

**Authors:** F. Paquin-Lefebvre, L. Kushnireva, Z. Xu, S. Kubler, T. Feofilaktova, S. Laughlin, N. Rouach, E. Korkotian, D. Holcman

**Affiliations:** Group of Data Modeling, Computational Biology and Predictive Medicine, École Normale Supérieure, Université PSL, Paris, France; Department of Immunology and Regenerative BiologyNeuroscience, Weizmann Institute of Science, Rehovot, Israel; Cambridge University and Churchill College, CB3 0DS, United Kingdom; Department of Biology, Perm State University, Perm, Russia

## Abstract

Synaptic activity imposes acute energy demands, especially for restoring ionic gradients via pumps and exchangers that require ATP. While mitochondria are positioned near dendritic spines to meet this demand, how ATP is produced and delivered with spatial precision remains unclear. Here, using high-resolution calcium and ATP imaging, immunocytochemistry, and computational modeling, we demonstrate that synaptic input—but not back-propagating action potentials (bAPs)—triggers on-demand mitochondrial ATP production. This occurs only in spines containing a spine apparatus (SA), where calcium-induced calcium release (CICR) activates mitochondrial calcium uniporters (MCUs), initiating ATP synthesis. We show that ATP delivery is spatially constrained to mitochondrial regions facing the spine base, where ATP-synthase is enriched. Importantly, ATP produced elsewhere on the mitochondrial surface tends to diffuse into the dendrite. We further demonstrate that the delivery of ATP to the spine head is geometrically optimized: an intermediate spine neck length maximizes delivery efficiency. Mathematical modeling and simulations revealb that the time scale for ATP to reach and refill all head-localized exchangers is on the order of hundreds of milliseconds—fast enough to meet local metabolic needs. The present findings establish a mechanism in which nanoscale calcium signaling and mitochondrial architecture together ensure rapid, spatially targeted ATP delivery, tightly coupled to synaptic activity.

## 1 Introduction

Fast cellular responses, such as synaptic transmission, typically occur on the millisecond timescale and incur little immediate energetic cost. However, the metabolic burden arises during repolarization, when cells must restore ionic gradients, insure vesicular trafficking and maintain membrane potential, energy intensive processes requiring ATP hydrolysis by pumps, exchangers or microtubule energy consumption [1, 2]. This delayed “energetic payback” has been well-documented in sensory systems, including photoreceptors and hair cells [3], and accounts for 60–80% of the energy consumed by neural signaling [1]. Astrocytes play also a fundamental role in regulating metabolism and providing substrate for neurons [4, 5, 6]

We investigate here the conditions under which ATP is produced in response to synaptic transmission. In neurons, ATP is primarily synthesized by mitochondria forming isolated or organelle networks typically fragmented and distributed throughout dendrites, with many positioned near the base of dendritic spines [7, 8]. Although this localization supports nearby energy demand, the geometry of the spine, particularly the narrow spine neck, poses potential barriers [9, 10, 11, 12] to electrical and chemical exchange, such as calcium signaling that can initiate ATP production and diffusion into the spine head. Moreover, mitochondrial dynamics and morphology can be regulated by synaptic activity through calcium-dependent fission and fusion events [13, 14].

The spine neck shapes calcium flow from the spine head to the dendrite [15, 16, 17, 18, 19]. This regulation becomes particularly relevant in the presence of a spine apparatus (SA)—a specialized endoplasmic reticulum (ER) structure enriched in certain spines—which amplifies calcium signals via calcium-induced calcium release (CICR) [20, 21, 22, 23, 24, 25]. CICR is initiated by synaptic calcium transients that activate ryanodine receptors at the base of the spine [23, 26, 27]. These amplified calcium signals can be locally transmitted to mitochondria via membrane-associated ER-mitochondria contact sites (MAMs), where calcium uptake through the mitochondrial voltage-dependent anion channel (VDAC on the outer membrane) and calcium uniporter (MCU on the inner membrane) is regulated by PDZD8 [28, 29]. Disruption of this interface impairs calcium homeostasis and mitochondrial signaling.

Under physiological conditions, ATP concentrations in spines are estimated at 1mM or less (0.41 *±* 0.01*mM* ). Given a typical spine volume of approximately 1 *µ*m^3^, this corresponds to *∼*3 *−*6 *×* 10^5^ ATP molecules [30]. This number appears disproportionate to the relatively low number of ATP-consuming molecules such as ion pumps and actin-remodeling proteins [31]. After a synaptic event, thousands of Na^+^ and Ca^2+^ ions enter the spine within milliseconds. For instance, a single millisecond event may introduce *∼* 6000 Na^+^ ions and *∼* 600 Ca^2+^ ions. To restore ionic gradients, ATP-driven pumps are engaged, each consuming one ATP molecule to transport three Na^+^ ions versus two in taken K+ ions [32] or exchanging one ATP per two Ca^2+^ ions [33, 34, 31]. To clear Na^+^ alone, approximately 30 pumps would be required, consuming around 2000 ATP molecules over one second. This discrepancy raises a central question: is the large steady-state ATP pool maintained continuously, or is ATP synthesized on demand to meet transient energy needs?

In this study, we focus on the mechanisms and constraints of on-demand mitochondrial ATP production following synaptic activity [6]. Using a combination of calcium imaging, mathematical modeling, and live cell confocal microscopy, we examine how synaptic inputs and back-propagating action potentials (bAPs) trigger calcium transients, ATP synthesis and delivery in spines with and without a spine apparatus. We quantify the correlation between cytoplasmic and mitochondrial calcium signals and resulting ATP production, and we model how nanoscale molecular spine architecture controls MCU activation and ATP delivery. Our results reveal that ATP synthesis is selectively triggered by synaptic input—but not by bAPs—and that efficient delivery into the spine head depends on the precise spatial mitochondrial membrane molecular component organization including MCU and ATP synthase that are actually facing the spine neck only. We further show that ATP produced outside of the spine base diffuses inefficiently and is largely lost into the dendritic shaft. Finally, based on diffusion modeling, we compute the timescale of ATP replenishment in the spine head and explore the role of the spine geometry and consumer density, providing a mechanistic understanding of local energy production during synaptic activity.

## 2 Results

### 2.1 Synaptic but not bAPs induce on-demand mitochondrial ATP production

To determine whether ATP production is triggered immediately following synaptic input, we simultaneously monitored calcium and ATP dynamics in dendrites and dendritic spines. In response to a synaptic event (*t* = 0), calcium concentrations peaked at approximately 120 ms in the spine head and 160 ms at the spine base in spines containing a spine apparatus (SA), as identified by synaptopodin labeling (Fig. 1A, top row), consistent with previous reports [35]. In spines lacking a SA, calcium peaks were delayed, reaching a maximum at 130 ms in the head and 210 ms at the base, and differed notably from baseline activity measured 5 *µ*m away (Fig. 1A–B).

**Figure 1.**
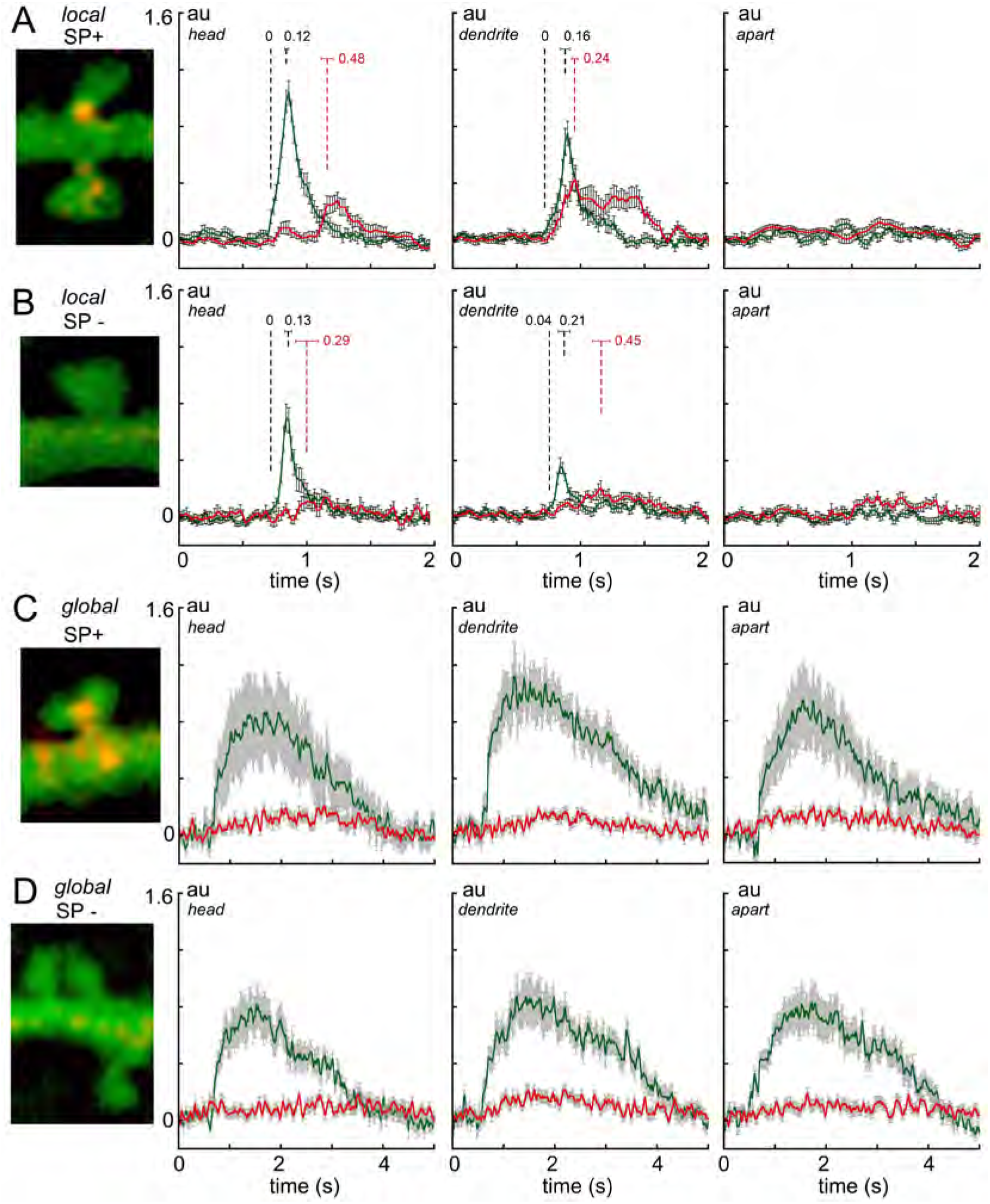
ATP levels following local and global spontaneous calcium events. **(A)** At 8 DIV after plating, hippocampal cultures were transfected with plasmids encoding: synaptopodin protein (SP-mCherry) the marker of spine apparatus/local endoplasmic reticulum and FRET-based ATP biosensor, ATeam (pcDNA-AT1.03). In about 17-23DIV weeks after plating, cultures were loaded with intracellular calcium sensors X-Rhod-1AM or Fluo-2AM, at room temperature during 30/60 min and imaged on Zeiss-880 confocal microscope, using 32-channel GaAsP detector and lambda-mode for accurate spectrum separation based on the pre-trained confocal system. Förster resonance energy transfer (FRET) was used for ATP-signal detection, presented as ΔF/Fs, red traces (A-D). Spontaneous calcium events are presented as ΔF/Fs, green traces **(A-D)**. Dendritic spines were classified into morphology groups as mushroom or stubby. Only the data for mushroom spines are presented in the figure. For stubby spines, see SI figure 1. Spines were divided into synaptopodin positive (SP+, A&C, red-yellow single or plural puncta) or negative (SP-, B&D). Global or local spontaneous calcium events were robustly specified using two criteria: duration of events (half decay within about 200 ms for local and about 2 s for global) and the presence of calcium transients at apart location within 5 *µ* m from the spine base, present during the global events (presumably back-propagating action potentials, C&D, right panels) and absent in local synaptic events (panels A&B, right panels). Signal detection was performed at the rate of 40ms, from 3 regions of interest (ROI) of equal size: spine head (left graph panels, A-D), dendrite (middle panels, A-D) or 5*µ*m apart (right panels, A-D).

Concurrently, ATP fluorescence signals revealed an initial rise in the spine head that closely followed the calcium transient, with a peak at 240 ms, followed by a plateau lasting several hundred milliseconds. In SA-containing spines, ATP levels in the dendrite peaked at 480 ms and subsequently rose in the spine head, indicating a delayed but coordinated energy delivery mechanism. In contrast, spines lacking a SA displayed no significant ATP production. In these spines, ATP level had a maximum at 290 ms in the head and 450 ms in the dendrite, with fluctuations comparable to the 5 *µ*m reference region, indicating that these signals likely represent background metabolic activity rather than specific on-demand ATP synthesis.

We next examined ATP dynamics following a back-propagating action potential (bAP). BAPs induced transient calcium elevations in both the spine head and dendrite, regardless of the presence of an SA (Fig. 1C–D). However, these calcium transients were not associated with any significant ATP increase, clearly distinguishing the response from that observed after synaptic input. We report similar dynamics for stubby spines (Fig. S1)

In summary, synaptic calcium transients in spines containing a spine apparatus trigger ATP production in adjacent dendritic mitochondria, with delayed propagation of ATP into the spine head approximately 500 ms later. Spines lacking a SA do not produce ATP in response to synaptic input, suggesting that the SA is required for calcium amplification via CICR, consistent with our previous findings on fast calcium communication [35]. Notably, ATP production persists for approximately 1 s before declining. These results demonstrate that synaptic input—but not bAP—can selectively activate mitochondrial ATP synthesis in an SA-dependent manner. In the following sections, we will examine the structural and molecular requirements for this selective activation.

### 2.2 Ca^2+^ diffusion model reveals that differential buffering between spines and dendrites supports selective MCU activation by synaptic input but not bAP

To better characterize the mechanism underlying the differential activation of mitochondrial ATP synthesis by synaptic inputs versus bAPs, we extended our previous Ca^2+^ diffusion model in dendritic spines [23, 36]. We incorporate here the spine apparatus, ryanodine receptors (RyR), and SERCA pumps, with RyR being the key mediator of CICR with additionally mitochondria positioned at the spine base (Fig. S2). We added MCU channels only in the upper-half (Fig. 2A), and modeled a differential calcium buffering by introducing a threefold higher calmodulin buffer concentration in dendrites compared to spines [37] (we used 120 vs 1700 buffers in the spine head vs dendrite, respectively to account for the difference in volumes 1.45 *µm*^3^ vs 0.3 *µm*^3^).

**Figure 2.**
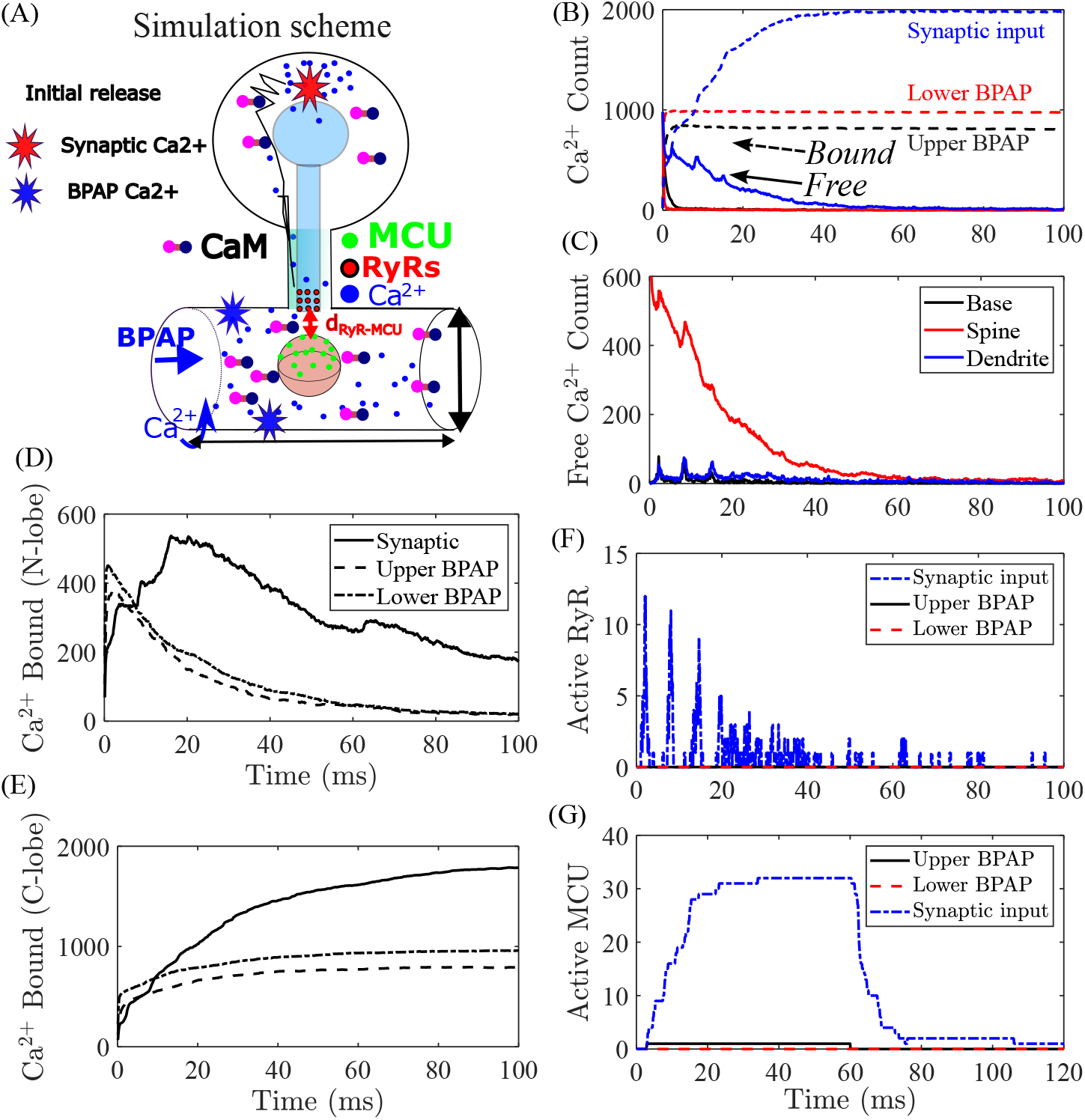
Simulation of Ca2+ dynamic during synaptic input versus back-propagating action potential and differential activation of MCU channel for a Mushroom spine. **(A)** Schematic of modeling MCU activation from synaptic input (top of the spine head, red star) vs bAP releasing calcium ion from the upper and lower part of the dendrite (blue stars). Calmodulin buffers consists of two lobes that bind ca2+ ions, limiting their diffusion. RyRs release Ca2+ via CICR and can lead to MCU opening (located on the upper part of the mitochondria). **(B)** Number of free (solid) and bounded (dashed) Ca2+ ions trigged by a synaptic input (blue), upper and lower bAP (black and red). **( C-D)** Number of Ca2+ bounded by the N- and C-lobe for the three types of calcium input. **(E)** Transient distribution of Ca2+ ions for synaptic input in the spine (red) and dentrite (blue) as well as the base of the spine (black). **(F-G)** Evolution of the numbers of active MCU and RyR for the three types of calcium input.

To compare how synaptic input and bAPs affect calcium signaling and mitochondrial activation in mushroom-shaped dendritic spines, we conducted stochastic simulations of calcium dynamics (see Method), buffering, and mitochondria MCU activation (Fig. 2A). We simulate synaptic input by introducing 1000 Ca^2+^ ions at the top of the spine head. In spines containing a SA and MCU channels on the mitochondrion located *∼*50 nm away, over a short period of less than 10ms, this input was sufficient to initiate various calcium effect: in response to synaptic input, free calcium peaks sharply within the first few milliseconds and is rapidly bound by intracellular buffers and targets, reaching a total bound calcium count of *∼* 2000 ions (Fig. 2B). In contrast, bAP (where Ca2+ in the dendrite initiate at blue stars in Fig. 2A) leads to weaker calcium signals, especially for calcium originating deeper in the dendrite, with lower binding saturation. Our simulations allow a spatial decomposition of free calcium and revealed a strong head-localized gradient for synaptic input, consistent with efficient local confinement of calcium (Fig. 2C). BAP-induced calcium, however, remains largely confined to the dendrite and does not significantly invade the spine head. We also separately analyzed calmodulin activation. Calcium binding to the N-lobe (fast kinetics, Fig. 2D) occurs rapidly and transiently, whereas binding to the C-lobe (Fig. 2E) accumulates over time. Both lobes show higher calcium occupancy in response to synaptic input compared to bAP.

We next examined RyR activation over time. Synaptic input leads to rapid and transient RyR opening within 5–10 ms, supporting CICR, while bAP-induced calcium failed to reliably trigger RyR opening, especially when calcium was released deeper in the dendrite (Fig. 2F). Finally, we quantified MCU activation (requiring three calcium ions to arrive within 20 ms). Synaptic input triggered (Fig. 2G) robust and sustained MCU opening (peaking at over 30 active MCU) probably due to the large amount of Ca2+ confined in the nano-domain between the SA and the mitochondria, whereas bAP failed to activate MCU altogether or activated it only weakly and transiently when calcium originated near the spine base.

In total, synaptic inputs activate more than half the number of MCU channels present in the upper-mitochondria (Fig. 2D). We run similar simulations in stubby spines (SI Fig. S3): while capable of initiating mitochondrial ATP production, they exhibit a reduced spatial selectivity due to their structural constraints. This suggests that mitochondrial activation in stubby spines is less efficient or temporally precise than in mature mushroom spines, thus providing an explanation for the differences we reported in Fig. 1. These simulations show that the structural arrangement of calcium sources, RyRs, and mitochondria permits selective activation of MCU and probably later on ATP synthesis pathways in response to synaptic input, but not bAP. This functional asymmetry is driven by local calcium amplification, spatial proximity, and kinetic dynamics at the molecular level.

We further evaluated the role of ER-SA distance when varying it from 50 to 200nm (Fig. S4A-B), we computed the peak (maximum) of RyR and MCU activated and found a clear decay when increasing the distance above 50nm. RyR activation (Fig. S4B) was less sensitive to this spatial separation, showing that the ER-mitochondria nanodomain does not confined Ca2+ enough to activated MCU. BAP failed to activate MCU or RyR channels, in contrast to synaptic inputs (Fig. S4 C-D).

Interestingly, to assess how intracellular buffering modulates these dynamics, we first increased the calcium buffer concentration in the spine head to match the dendritic level (x3 increase), this strongly attenuated free calcium levels, which dropped below 10 MCU within 70ms post-stimulus. Conversely, decreasing dendritic buffering (Fig. S4E black curves) led to a drastic increased MCU activation and restored a mitochondrial engagement similar to synaptic inputs. These results show that intracellular buffering landscapes critically regulate CICR amplitude and mitochondrial recruitment. Thus calcium buffering is key to prevent bAP to activate MCU. Finally, we examined how the spatial distribution of MCU on the mitochondrial surface influences activation efficiency. We compared upper-only MCU localization (facing the ER) with uniform and lower-only distributions. Uniform distribution yielded 50% fewer activated MCU channels compared to the upper-only condition (Fig. S4G), while lower-localized MCU channels remained largely inactive. Notably, RyR activation remained unchanged across conditions (Fig. S4H). This result predicts that the spatial orientation of MCU relative to the ER is key to maximize the activation likelihood.

Collectively, these numerical simulations reveal that successful MCU activation is not solely a function of calcium release but depends critically on spatial coupling, buffer environment, and mitochondrial molecular topography. At this stage, we conclude that the buffering capacity, MCU organization and nanometer organization between the spine apparatus and mitochondria regulate MCU calcium activation selectively to synaptic inputs but not bAP.

### 2.3 Optimal ATP delivery occurs at an intermediate spine neck length

We next study how Ca2+ signal triggers mitochondria ATP production and whether the efficiency of ATP delivery to the spine head depends on the spine morphology such as the neck length *L*_neck_ or spine head radius *R*_Head_. As ATP-consumers such as ion exchangers and actin-remodeling motors are distributed across the dendritic spine, we though that the spine geometry could have an influence on ATP consumptions. To assess this, we quantified the peak ATP concentration 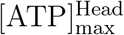 in the spine head, across spines for various *L*_neck_. Surprisingly, we found a non-monotonic relation for [ATP]^Head^ vs *L*_neck_ with a maximum at an optimal neck length 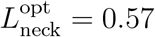 *µ*m (*p* = 3.4*×*10^*−*3^, *n* = 30 spines, as shown in Fig. 3A). This result suggests that longer spine necks may result in ATP loss—either through consumption during diffusion or through a less efficient production mechanism at the base.

**Figure 3.**
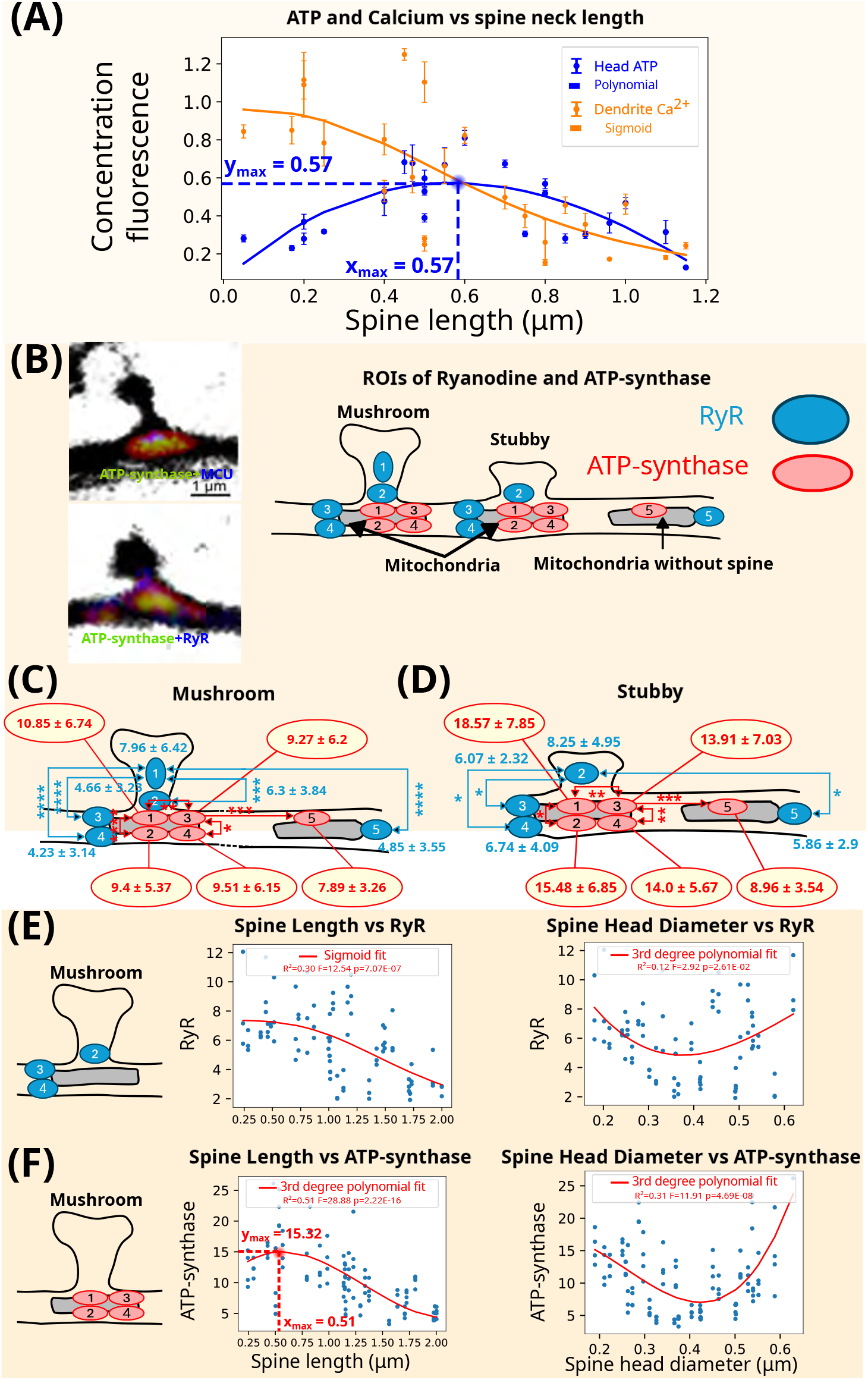
Distribution and Correlation of Ryanodine Receptors (RyR) and ATP-Synthase in Mitochondria at Dendritic Spine base. **(A)** ATP in the spine head and calcium concentrations in the dendrite versus spine length. Calcium decrease is fitted by a sigmoid function equation 9 with parameters 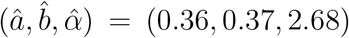, with (*R*^2^, *F*_*stat*_, *p*) = (0.55, 8.22, 9.27 *×* 10^*−*4^). ATP in spine head is fitted with a degree 3 polynomial function (eq.8) with parameters 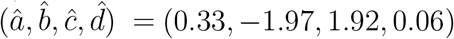 with (*R*^2^, *F*_*stat*_, *p*) = (0.55, 5.72, 3.4 *×* 10^*−*3^). The maximum ATP in spine head is obtained for a length *L*_*max*_ = 0.57*µm*. **(B) Left:** Immunostaining of ATP-synthase + MCU vs ATP-synthase+ RyR: with morphology (BFP) in Black mitochondria morphology in Red. (MT DsRed) and ATP-synthase (antibody) in Green. Blue: upper image is MCU while lower right image represent RyR. **Right:** Segmentation of mitochondria near mushroom and stubby dendritic spines in Regions of Interest (ROIs) for RyR (blue) and ATP-synthase (red). **(C)** Mean and standard deviation (*µ, σ*) of the RyR and ATP-synthase distributions in ROIs for mushroom spines. The statistical significance of the decrease of the ATP-synthase and RyR distributions with the distance from spine head is computed using a paired one-tailed Wilcoxon signed-rank test (n.s: *p >* 0.05; *: *p <* 0.05; **: *p <* 0.01; ***: *p <* 0.001; ****: *p <* 0.0001). Region *R*_1_ is the reference distribution both for RyR and ATP-synthase. **(D)** Mean and standard deviation (*µ, σ*) of RyR and ATP-synthase in mitochondrial and dendritic regions. Statistical significance of the decrease of the ATP-synthase and RyR distribution with distance from spine head uses the paired one-tailed Wilcoxon described above. Region *R*_2_ for RyR and *R*_1_ for ATP-synthase are used as reference distributions. **(E)** Correlation between RyR vs spine length and spine length diameter in mushroom spines uses the aggregated ROIs *R*_2_, *R*_3_, *R*_4_. RyR decrease is fitted by a sigmoid function equation 9 with parameters 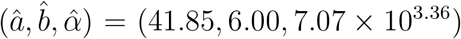 with (*R*^2^, *F*_*stat*_, *p*) = (0.30, 12.54, 7.07 *×* 10^*−*7^). RyR against spine head diameter is fitted by a degree 3 polynomial (eq. 9) with parameters 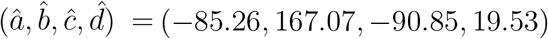 with (*R*^2^, *F*_*stat*_, *p*) = (0.12, 2.92, 2.61 *×* 10^*−*2^). **(F)** ATP vs spine length and spine head diameter in mushroom spines obtained by aggregating ROIS *R*_1_, *R*_2_, *R*_3_, *R*_4_. ATP-synthase versus spine length is fitted by a degree 3 polynomial function (eq. 9) with parameters 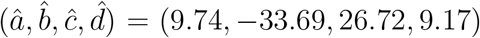 and (*R*^2^, *F*_*stat*_, *p*) = (0.51, 28.88, 2.22 *×* 10^*−*16^). ATP-synthase versus spine head diameter is fitted by a degree 3 polynomial (eq. 9) with parameters 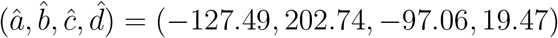 and (*R*^2^, *F*_*stat*_, *p*) = (0.31, 11.91, 4.69 *×* 10^*−*8^).

To test whether this optimal ATP profile could be explained by Ca^2+^ signaling, we plotted the calcium levels in the spine head vs *L*_neck_. Instead of a maximum, we observed a monotonic decrease in calcium concentration with increasing neck length (*p* = 9.27 *×* 10^*−*4^), indicating that the ATP delivery optimum does not directly result from an optimal calcium dynamic. In addition, ATP could be consumed in the neck. At this stage, we conclude that for small spine neck, Ca2+ appears at the base, leading to a small amount ATP in the head, possibly due to a leaking effect. For long neck, Ca2+ becomes small, which leads to less produced ATP. An optimal spine length could fulfill a role for optimal spine computing [17] and this optimum may arise from enhanced ATP synthesis along the spine neck that we shall explore below.

### 2.4 Spatial organization of RyR, ATP-synthase and MCU correlates with optimal ATP delivery and selective mitochondrial activation

To further investigate the molecular basis of selective ATP production by synaptic input and the optimal ATP concentration in the spine head, we examined simultaneously the spatial distribution of RyR which mediates calcium amplification, and mitochondrial ATP-synthase, which exchanges ADP to ATP at the mitochondrial surface.

To quantify the co-localization between RyR and ATP-synthase, we subdivided the spine and adjacent dendrite into five regions of interest (ROIs): R1–R2 encompassed the spine head and neck; R3–R4 spanned the peri-synaptic dendrite; and R5 served as a distal dendritic reference (Fig. 3B). Immuno-staining revealed that RyR expression was enriched in R1 and R2 (Fig. 3C), with a clear preference for mushroom spines over stubby spines, consistent with prior work [38]. In contrast, RyR expression was markedly lower in dendritic ROIs (R3–R5).

Similarly, we analyzed the distribution of ATP-synthase. We found that ATP-synthase was enriched at the mitochondrial membrane surface facing the spine base (R2), and this enrichment was more pronounced in stubby spines compared to mushroom spines (Fig. 3D). This suggests that mitochondrial positioning and ATP-synthase orientation are optimized for directing ATP export into the spine head, potentially minimizing loss into the dendrite. To explore how these protein distributions relate to spine morphology, we correlated RyR and ATP-synthase expression with spine neck length *L*_neck_ and head diameter *D*_Head_. For mushroom spines, RyR expression in R1–R2 decreased with increasing *L*_neck_, suggesting reduced CICR capability in longer spines. Interestingly, RyR expression exhibited a non-monotonic relationship with *D*_Head_, showing a local minimum that remains difficult to interpret (Fig. 3E).

Pooling ATP-synthase expression across R1–R4, we found that its expression peaked at a neck length of *L*_neck_ = 0.52 *µ*m (Fig. 3F), remarkably close to the optimal length for ATP delivery identified earlier. This spatial correlation suggests that optimal mitochondrial ATP delivery is achieved not only through calcium signaling, but through the spatial arrangement and expression levels of ATP-synthase. However, the mechanism by which this ATP-synthase localization is established and maintained at the molecular level remains an open question.

To further investigate the nanometer molecular organization of input (Ca2+)–output (ATP), we explore the correlative distribution of MCU versus ATP-synthase. Our Ca2+ simulations described above predicted that MCU should be preferentially located toward the entrance of the spine to favor the reception calcium generated by the RyR and thus should be co-localized with high ATP-synthase expression. Such colocalization would also define a memory path of release ATP toward the preferential direction of the input calcium signal. To test this hypothesis, we thus perform immunocytochemistry with both MCU and ATP-synthase (Fig. S5A-C and S6), where we found a significant increase of the co-expression of both in the region just below the spine (region 1) with a significant gap for mushroom spines with the adjacent regions (2-4) or the comparison region 5 (isolated mitochondria). This trends remains for stubby spine, but less pronounced. We also found a positive correlation (linear regression between ATP-synthase and MCU) and an optimal MCU expression versus spine length for *l* = 0.8*µm* (see Fig. S5 for other correlation with the head diameter).

To further investigate the intermingling of ATP-synthase and MCU within mitochondria, we first computed the shortest mean distance between an ATP-synthase and MCU molecules and found a distance of 150nm (Fig.S6). Second to quantify the degree of intermingling between the two population, we identified the convex hull of the high-intensity pixels for both populations inside each mitochondrial mask and computed the intersection over Union (IoU) metric 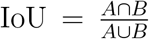, where *A* and *B* are the surface of the convex hulls corresponding to ATP-synthase and MCU molecules, respectively. We found an IoU of 50% (Fig. S6), corresponding to a high covering between the two populations. Similarly, we estimated the spatial coupling from the mitochondrial volume overlap 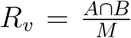, where *M* is the total mitochondrial volume, leading to a similar result 54%. We conclude that ATP-synthase and MCU have a high degree spatial correlation and represent two well mixed clusters of molecules, confirming the hypothesis of a molecular memory re-arrangement or trace between Ca2+ entrance signal and optimally exported ATP molecules toward the spine head.

### 2.5 Deciphering on-demand ATP production and delivery

We quantify here ATP concentration transported in the head, as measured in Fig. 1A). By fitting the initial slope of ATP production (Fig. 4A-B), we obtain a production rate of *κ*_*ATP−production*_ = 2Δ*F/Fs*. Assuming a range of 1-5mMol of total ATP in neurons, we found that the maximum corresponds to 6% increase and thus in 1*µm*^3^, we have for 1 mMol 6 *\** 10^5^. We conclude that excess of ATP produced at the maximum triggered by synaptic input is 127,570 molecules (peak below spine in mushroom), which is reduced to 81,425 for stubby, while in the head it is 83,581 (mushroom) and 61,635 (stubby). We further extract the production *κ*_*ATP−prod*_ and consumption *κ*_*ATP−cons*_ rates by fitting the initial and decay slopes for the difference of the ATP at the base and in the head over n=18 experiments (Fig. S7). We obtain for the production and the consumption rate *α*_*P*_ = 542, 759 ATPs/s (resp. *α*_*C*_ = *−*338, 209) for Mushroom and *α*_*P*_ = 455, 278ATPs/s (resp. *α*_*C*_ = *−* 53, 341) for stubby spine.

**Figure 4.**
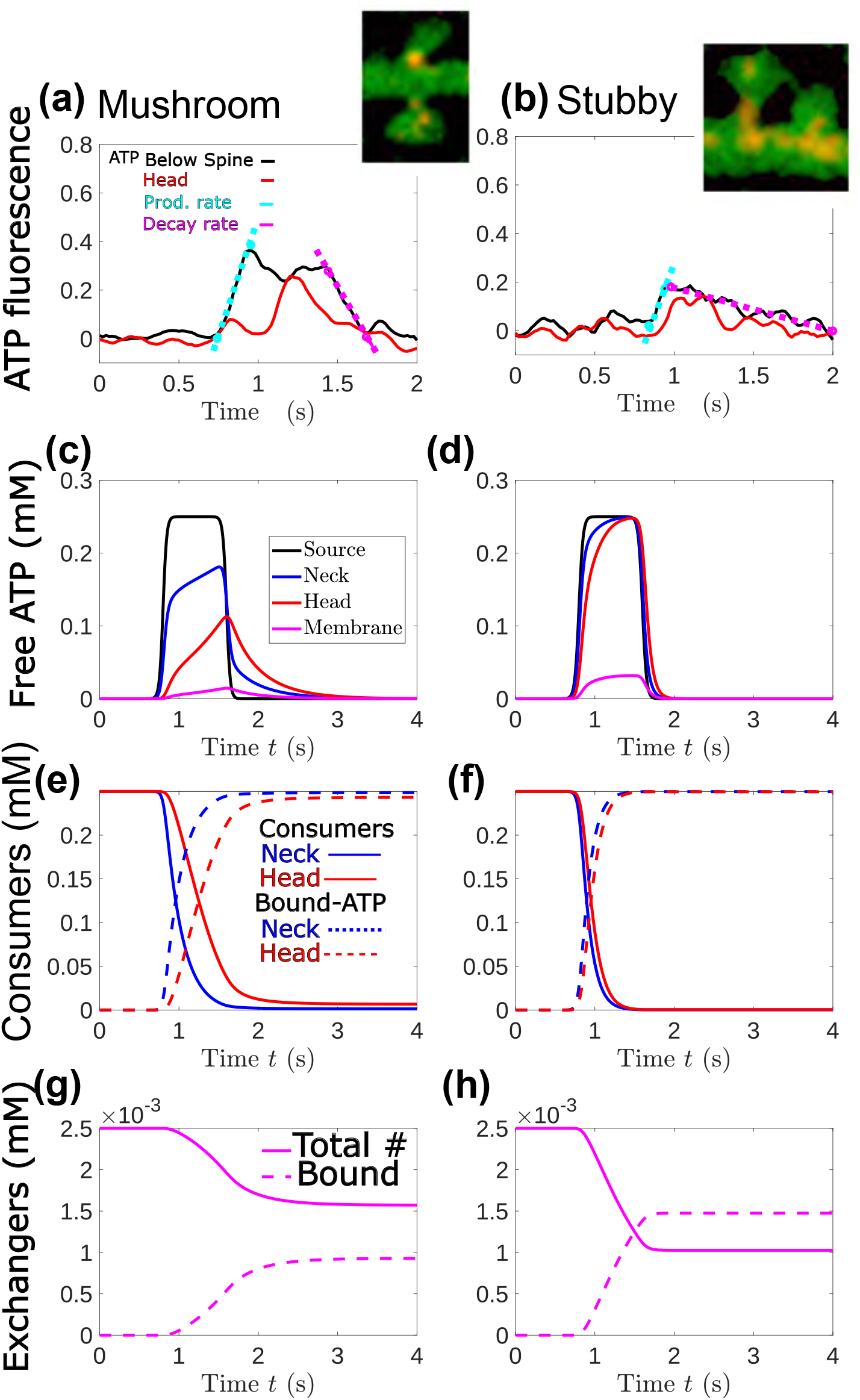
Experimental vs modeled ATP dynamic in mushroom and stubby dendritic spines. **(A–B)** ATP fluorescence changes measured in mushroom (A) and stubby (B) spines using FRET-based biosensors. Traces show ATP levels below the spine (black) and in the spine head (red). Cyan and magenta dots indicate fitted rising and decay phases, respectively. Insets: representative spine morphologies from confocal imaging. **(C–D)** Modeled dynamic of free ATP concentration over time at four representative compartments: ATP production source-spine base-(black), spine neck (blue), spine head (red), and membrane-bound exchanger region (magenta). Results in (C) and (D) are obtained with COMSOL Multiphysics [58]. **(E–F)** Evolution of cytosolic ATP consumers (solid lines) and ATP-bound consumer complexes (dashed lines) in the neck (blue) and spine head (red). **(G–H)** Binding dynamic at the head spine membrane: total number of free exchangers (solid magenta) and ATP-bound exchangers (dashed magenta). Mushroom spines (left panels) exhibit faster and more localized ATP accumulation and exchanger refilling compared to stubby spines (right panels), as predicted by the compartmental reaction–diffusion model (see SI for method details and associated kymographs, Fig. S12).

By subtracting the transported ATP from the base (fast diffusion) with the one to the head, we reported two phases: a first one lasting hundreds of ms (first bump in Fig. S7) followed by a second bump. These bumps suggest two production-consumption phases, well marked for mushroom but less pronounced for stubby spines. To further characterize these phases, we developed a reaction–diffusion model (Method) that accounts for ATP diffusion, irreversible binding to immobile cytosolic consumers, and interaction with membrane-bound exchangers located in the spine head.

Here ATP binds to consumer and exchangers according to the chemical equations

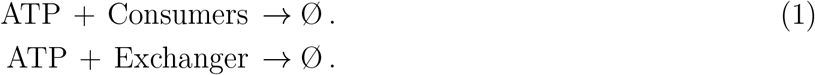

Using the extracted production and consumption rates, we accounted for the first production phase, but not the second one (Fig. 4C-D). This suggests an extra-mechanism that should be triggered when ATP concentration reaches a high concentration, such as local protein synthesis. When we added in the model the presence of exchanger located on the spine head surface, we found that they are activated in a time scale of hundreds of ms, due to the first consumption of ATP by internal ATP-consumers (Fig.4E-H) after a first phase where ATP is directly hydrolyzed by internal consumers, (Fig.4E-H), exchangers are also refilled quickly (see also our other variant modeling Fig. S8-S9). To conclude, the ATP time scale in the spine is driven by the balance between the mitochondria production rate and the number of consumer with their hydrolysis rate. In that case, both diffusion and consumer rate contribute to the time for exchanger to be activated of the order of hundreds of ms. Finally, our model predicts in stubby spines, the absence of a constricting neck facilitates ATP spread from the mitochondria to the head compartment (no diffusion gradient), resulting in higher exchanger ATP-recovery. Simulations predict that 59% of exchangers in stubby spines will recover a functional state following ATP biding, compared to only 37% in mushroom spines (Fig. 4), underscoring the influence of spine geometry on metabolic coupling at excitatory synapses.

### 2.6 Mitochondrial ATP distribution between dendrite and spine

To quantify the spatial fate of ATP molecules synthesized by mitochondria at the base of dendritic spines, we modeled the partition of ATP between the dendrite and the spine head. ATP must reach ATP-dependent molecular targets such as Na^+^/K^+^-ATPases, Ca^2+^-ATPases (e.g., PMCA, SERCA), and exchangers embedded in the spine head membrane. However, during its transit, ATP may also be consumed by cytosolic processes such as actin remodeling, receptor trafficking, and initiation of local protein synthesis. These competing pathways shape the effective delivery of ATP to postsynaptic targets. We first characterize this spatial distribution by estimating the probability that an ATP molecule reaches the spine head exchangers versus escaping into the adjacent dendritic shaft. This splitting probability depends critically on spine geometry and the initial release position of ATP.

The probability *P*_Exchangers_(0) that an ATP molecule released at the base of the spine enters the head (in the absence of cytosol consumers) *≈*18%, indicating that fewer than 1 in 5 ATP molecules successfully reaches the spine head exchanger targets (Fig. S6-S7) if released directly at the base of the neck. This result aligns with our stochastic simulations (see Fig. 5, initial release at *z* = 2000 nm). To assess how spine morphology affects this probability, we applied the model to the geometric parameters of mushroom and stubby spines (Table 1). Differences in neck length and mitochondria-to-spine distance have a pronounced effect on ATP capture efficiency. Stubby spines—with shorter necks and closer mitochondrial proximity—exhibit a significantly higher ATP delivery probability (factor 2: from 0.2 vs 0.1, Fig. S8-S9) than mushroom spines. These morphological features likely contribute to the greater metabolic efficiency observed in stubby spines. To better characterize the difference in ATP production between stubby versus Mushroom spine, we also analyzed the distances between mitochondria and in each spine type. The mean distance from mitochondria to mushroom spines was 137 *±* 137*nm*, indicating a highly localized mitochondrial positioning with relatively low variability (Table 1). In contrast, the mean distance to stubby spines was significantly larger at 353 *±* 219*nm*, reflecting greater spatial variability and overall displacement. These findings suggest that mitochondria are preferentially and more consistently positioned near mushroom spines, facilitating more efficient ATP delivery to support synaptic activity in these mature structures.

**Figure 5.**
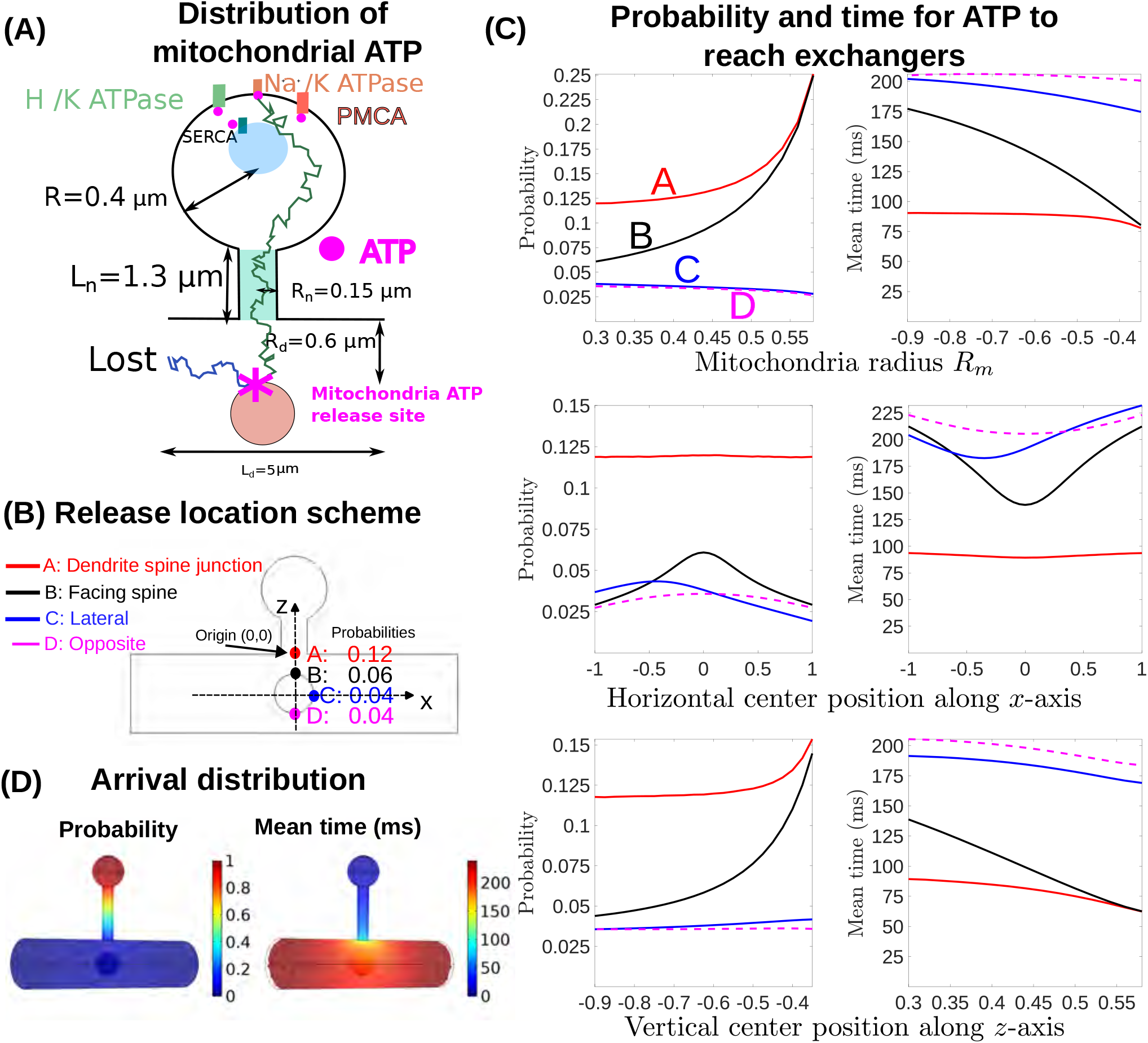
Efficiency of ATP arrival at spine head exchangers for mushroom spine. Probability distribution and mean arrival time of an ATP molecule vs release site within the dendritic spine. **(A).**Scheme of an ATP reaching various either an exchangers (including Na^+^/K^+^-ATPase, H^+^/K^+^-ATPase, SERCA, and PMCA exchangers) or bing lost in the dendrite vs initial position on the mitochondria. Parameters are used from mushroom dendritic spines. **(B)** Release locations relative to the spine plasma membrane, **(C)** Arrival probability and mean arrival time vs release position (e.g., base, lateral, or head) and on mitochondrial geometry (mitochondrial radius and position with respect to the spine). **(D)** Probability and times for ATP to reach exchangers located on the spine surface, emphasizing how spatial release influences energy delivery.

**Figure 6.**
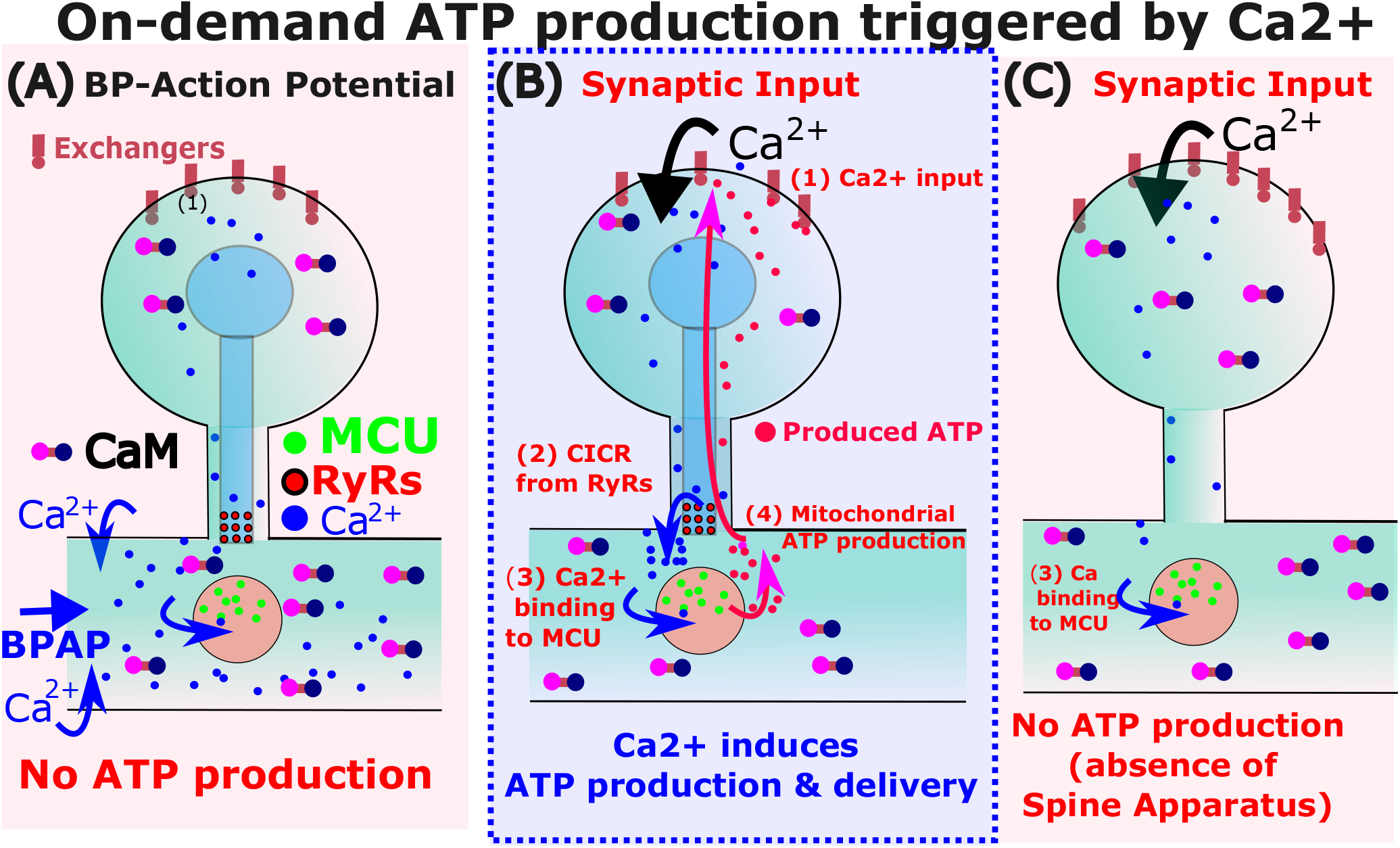
Summary of the on-demand ATP production at synapses. **(A)** BAP leads to calcium release from voltage channels, but no ATP production at mitochondria sites. **(B)** Synaptic input triggers calcium entrance in the spine head followed by inducing Calcium-Induce-Calcium-Release (CICR) from the Spine-Apparatus at the base of the spine. This result in high calcium at nanodomain, leading to the activation of MCU channel located on the mitochondria. These events are associated to ATP production and delivery to ATP-consumers located spine head. **(C)** Synaptic inputs in spines with no Spine Apparatus cannot trigger CICR and thus no ATPs are produced.

**Table 1.** Morphological parameters of mushroom and stubby spines.

| Parameter | Mushroom Spines | Stubby Spines |
| --- | --- | --- |
| Depth of mitochondria from spine base | $0.08 \pm 0.036 \mu\text{m}$ | $0.28 \pm 0.084 \mu\text{m}$ |
| Mitochondrial diameter | $0.76 \pm 0.039 \mu\text{m}$ | $0.67 \pm 0.087 \mu\text{m}$ |
| Mitochondrial length | $3.10 \pm 0.254 \mu\text{m}$ | $5.47 \pm 0.749 \mu\text{m}$ |
| Dendrite diameter | $1.17 \pm 0.070 \mu\text{m}$ | $1.19 \pm 0.147 \mu\text{m}$ |
| Spine head diameter | $0.79 \pm 0.066 \mu\text{m}$ | $0.86 \pm 0.046 \mu\text{m}$ |
| Spine length | $1.29 \pm 0.083 \mu\text{m}$ | $0.52 \pm 0.037 \mu\text{m}$ |

**Table 2.** Parameters of the stochastic simulations and analytical calculations.

| Parameter | Symbol | Value(s) |
| --- | --- | --- |
| Time step | $\Delta t$ | $10^{-7}$ s |
| Diffusion coefficient | $D$ | $350 \mu m^2 s^{-1}$ [57] |
| Spine head radius | $R$ | $0.5 \mu m$ |
| Spine neck radius | $R_n$ | $0.15 \mu m$ |
| Spine neck length | $L_n$ | $0.5 \mu m$ |
| Dendrite radius | $R_d$ | $1 \mu m$ |
| Dendrite length | $L_d$ | $5 \mu m$ |
| Circular obstacle radius | $r_m$ | $0.1 - 0.9 \mu m$ |
| Number of absorbing targets | $n_r$ | $\approx 50$ |
| Circular target radius | $A$ | $10$ nm |

These differences in the distance between mitchondria and the spine neck entrance is sufficient to account for the difference in the ATP delivered in the spine head and probably the lower production rate as less calcium ions can activate MCU. Indeed, Accounting for positioning of ATP-synthases at various distances such as *z* = *−* 300*nm* and *z* = *−* 100*nm* for stubby and mushroom spines and accounting also for their difference in geometry, we found that delivered ATP to exchangers (SI section 10) is 5% and 10% respectively. The smaller probability for the mushroom spine is due to the longer neck. We conclude that the ATP efficiency for the two types of spine is of similar magnitude, but can also be influenced by the mitochondria geometry, as shown by stochastic simulations where all ATP molecules are placed instantaneously at the mitochondria surface (Fig. 5A). Here ATP molecules can diffuse and each trajectory can either stay inside the dendrite and then travel far away or enter the spine head where they can ultimately reach the exchangers located on the surface. When we consider *n*_*r*_ = 50 exchangers that can hydrolyzed ATP molecules (modeled as consuming each ATP upon arrival), we found that the probability of ATP molecules to reach exchangers is 12% when release on the mitochondria facing the spine neck, it decreases to 6% on the side (Fig. 5B), finally the percentage continue to decay to 4% for any release below the mitochondria. To conclude, release at the top of the mitochondria is crucial for spine replenishment (similar results are presented for stubby spines in Fig. S10-11).

To further explore the impact of the initial ATP release location on the fraction that enter into the spine head and bind to the exchangers vs staying in the dendrite, we simulated ATP molecules by first varied the initial position in the horizontal y-axis direction (Fig. 5C-D) and then along the central z-axis of the dendrite. Finally, to evaluate the impact of an initial release along the centered (for y=0) z-axis vertical of the initial ATP release location, (Fig. 5C-D), we varied the position from the lower part of the dendrite *z* = 0 up to the upper opening of the spine neck for *z* = 2500, leading large changes. We found that the ATP fraction reaching the spine surface is very sensitive to the size of the mitochondria (radius *R*_*m*_) and to a shift in both the horizontal and the vertical axis, leading to factor 10 differences (Fig. 5D) in the ATP-distribution. To conclude, optimal ATP delivery requires a refined located of mitochondria at the base of the spine, so that mitochondria ADP-ATP-exchangers located toward the entrance of the neck can provide a maximal ATP-influx.

Finally, we needed to quantify the time scale of ATP delivery. To do so, we define the time scale of exchanger refilling the time for ATP to travel from mitochondria surface to reach an ATP-exchanger site in the absence of other ATP consumers. This time (see formula 3) can be estimated using values for the head radius *R* = 0.5*µ*m, neck length *L*_*n*_ = 0.5*µ*m, neck radius *R*_*n*_ = 0.15*µ*m and exchanger target radius *A* = 0.01*µ*m and a cytoplasmic diffusion constant of *D* = 350*µ*m^2^*/*s. Accounts for crowded dendritic spine, where the ATP-diffusion coefficient is diminished (factor 20 [39]) leading to an effective diffusion *D*_*eff*_ *≈* 15*µm*^2^*/s*, we obtain an average time for one ATP molecule to bind to one exchanger of 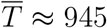 ms, Another quantification to estimate the time is to study the time for the last exchanger to be activated (formula 5, see method and SI Fig. 5) of the order of hundreds of milliseconds (Fig. S13). To conclude, the time scale of exchanger refilling by diffusion in a crowded dendritic spine is of the order of 1*s*. Such a time could be increase due to the production rate (Fig. 4) and the presence of a large amount of consumers.

## 3 Discussion

In this study, we uncover nanometer-scale organization and mechanism by which synaptic activity—specifically local excitatory input—elicits mitochondrial ATP production and targeted delivery to the dendritic spine head. This process is not constitutive but instead triggered on demand by localized calcium amplification via the spine apparatus (SA). The SA acts as a Ca2+ amplifier, allowing CICR through RyR channels to elevate calcium levels to the threshold required for mitochondrial calcium uniporter (MCU) activation. We further reported here a time scale ATP production of hundreds of milliseconds to one second, while ATP can be found in spine head in hundreds of millisecond after Ca2+ invades the spine head.

In contrast, bAPs, despite generating detectable calcium elevations, fail to activate MCU due to stronger dendritic buffering and the absence of activation of amplified Ca2+ at the ER-mitochondria contact. Our modeling and simulations confirm that mitochondrial ATP production is directionally biased: ATP generated at the mitochondrial surface facing the spine base reaches ATP exchangers with a splitting of 80% in the head within tens of milliseconds, whereas ATP synthesized elsewhere is largely lost to the dendrite. By quantifying ATP refilling timescales, identifying delivery optima, and demonstrating a critical dependence on spine architecture, we show here that the spatial nano-coupling architecture between calcium influx, RyR activation, MCU localization, and ATP synthase generates a functional bioenergetic synaptic microdomain.

### 3.1 Bimodal mitochondrial ATP response

Dendritic spines are supposed to maintain ATP at millimolar concentrations through continuous mitochondrial production [2, 40, 31]. However, this assumption is difficult to reconcile with the relatively low energetic cost of a single excitatory postsynaptic potential (EPSP), which should require only a few thousand ATP molecules—several orders of magnitude less than the presumed steady-state pool. Our findings support an alternative view: that ATP availability in spines follows a bimodal regime, with a low basal concentration sufficient for routine maintenance, and a rapid, spatially restricted burst of synthesis triggered by synaptic activity. This on-demand ATP production, observed only in mature spines equipped with a spine apparatus, enables localized mitochondrial responses via CICR and activation of the VDAC/MCU.

In spines lacking this capability, sodium influx following synaptic input cannot be effectively cleared by ATP-dependent exchangers and must instead dissipate through the spine neck. This could lead to altered voltage dynamic and potential disruption of local membrane properties. ATP scarcity in such compartments may also impair energetically demanding processes like AMPA receptor trafficking [41] and actin cytoskeleton remodeling, both essential for long-term synaptic maintenance.

Additionally, we report here a two-phase ATP consumption profile in mushroom spines. An initial rapid depletion phase reflects immediate consumption by high-affinity exchangers, while a second delayed decline occurs even as basal ATP levels near the spine base remain stable. This unexpected second-phase decay suggests the activation of a distinct class of ATP consumers or delayed recruitment of metabolic processes not accounted for by passive diffusion or extrusion through the neck. The origin and functional significance of this biphasic consumption warrant further investigation.

Finally, comparing spine morphologies, we observed that stubby spines (*L ≤* 0.5 *µ*m) exhibit faster ATP accumulation than mushroom spines (*L ∼*1 *µ*m), due to shorter diffusion paths and reduced buffering volume. The overall production time scale—approximately 1 second—is compatible with previously reported mitochondrial metabolic kinetics [42]. The characteristic time for ATP diffusion and binding to molecular consumers within the spine head was found to be on the order of hundreds of milliseconds. This timescale is consistent with estimates from diffusion theory, which models the binding kinetics of diffusing molecules in confined geometries [43]. To conclude, the present data analysis and our simulations predict that under optimal alignment between mitochondria and spine neck, all major ATP-consuming pumps such as the Na^+^/K^+^-ATPase can be refilled by at least one ATP molecule within hundreds of milliseconds, provided that ATP synthesis is triggered immediately following calcium activation. This rapid refueling capability highlight show spatial and temporal coupled energy production maintains synaptic function.

### 3.2 Mitochondrial Positioning at the Spine Base Enables Localized Calcium-Triggered ATP Production

The spatial positioning of mitochondria in close proximity to the base of dendritic spine necks plays a pivotal role in regulating both calcium-triggered ATP production and the efficient delivery of ATP into the spine head. The presence and precise localization of the spine apparatus (SA) is essential for calcium amplification through CICR [44, 35]. RyRs are specifically clustered at the base of the spine, enabling a local amplification of calcium signals that is sufficient to activate MCU. This is further supported by our current findings (Fig. 3), which show a strong spatial enrichment of MCU molecules in the nanodomain just beneath the RyR cluster.

Importantly, we previously reported a depletion of SERCA pumps in this basal region compared to the spine head, ensuring that calcium transients are not prematurely buffered and thus enabling robust mitochondrial activation. Any disruption of the SA–mitochondria alignment impairs this mechanism, as previously discussed [28, 29].

Our simulations (Fig. 5) reveal that MCU activation is highly sensitive to the distance between the mitochondrion and the RyR cluster: distances below 50 nm result in maximal MCU activation (Fig. 5), while distances greater than 300 nm, lead to a dramatic decline in calcium influx and mitochondrial responsiveness. Consequently, mushroom spines with mitochondria positioned 100 nm from the spine base are far more efficient in triggering on-demand ATP synthesis compared to morphologies with increased separation. Thus, nanoscale mitochondrial positioning is not only necessary but functionally optimized for rapid ATP generation following localized calcium release via the SA.

This precise spatial arrangement extends to ATP delivery. We show that ATP synthase complexes are enriched on mitochondrial membranes apposed to the spine base. This spatial bias ensures that newly synthesized ATP efficiently enters the spine neck and reaches local energy-demanding sites, including ion pumps such as the Na^+^/K^+^ exchanger. Misalignment of this mitochondrial output face leads to diffusion of ATP into the dendritic shaft, significantly reducing the metabolic support available to the synapse. The colocalization of MCU and ATP synthase within a constrained nanodomain forms a directional and functionally privileged pathway for ATP flow. This architecture acts as a form of “molecular memory path,” encoding the spatial origin of the calcium signal and coupling synaptic input with metabolic output.

Moreover, our data suggest that this nanoscale arrangement—comprising RyR, MCU, and ATP synthase—is essential for efficient calcium signaling between the SA and mitochondria, and for the targeted delivery of ATP to the spine head. Such an organization is not observed in mitochondria located away from spines, indicating a co-adaptive mechanism between dendritic spines and nearby mitochondria. This raises intriguing questions about the origin and maintenance of such sub-organelle architecture. We hypothesize that this reflects a form of synaptic-mitochondrial co-plasticity that may be spine-type specific, as suggested by differences observed between mushroom and stubby spines. Analogous nanoscale enrichment and compartmentalization have been previously described in the context of synaptic receptor organization. AMPA, NMDA, and glycine receptors are known to aggregate in nanodomains at the postsynaptic density, stabilized by specific scaffolding proteins [45, 41], and are aligned with presynaptic vesicle release zones via “nano-columns” [46, 47, 48]. Similarly, it is conceivable that transmembrane tethering proteins mediate the alignment between the SA and mitochondria. Candidates include PDZD8, which has been shown to colocalize with ER-resident FKBP8 and form functional tethers between organelles [29]. Such molecular scaffolds may serve as organizational hubs around which MCU, RyR, and ATP synthase complexes aggregate to support compartmentalized bioenergetics and calcium signaling at the single-synapse level.

### 3.3 How ATP is used in dendritic spines

We report here that up to 600,000 ATP molecules per second can be produced locally at dendritic mitochondria during the first few hundred milliseconds following synaptic stimulation (Fig. 4). This production, triggered by calcium-induced activation of the MCU, is part of an “on-demand” metabolic response finely tuned to local neuronal activity. Notably, this quantity vastly exceeds the number of ATP molecules needed to restore ionic gradients following a single synaptic event—on the order of a few thousand, primarily for Na^+^/K^+^ ATPases and Ca^2+^ pumps [1, 2]. The surplus ATP supports a broad range of critical processes within the dendritic spine. These include actin cytoskeleton remodeling required for spine motility and structural plasticity, recycling and trafficking of membrane proteins, endocytic and exocytic events, proteostasis, and maintenance of organelle function [49, 50, 51]. ATP is also consumed during protein turnover, phosphorylation cascades (e.g., via kinases like CaMKII), and vesicular trafficking—all processes known to be up-regulated following synaptic activity. Importantly, our data and simulations show that the spatial positioning of ATP-consuming elements relative to the site of mitochondrial ATP synthesis significantly impacts the temporal dynamic of energy delivery. ATP must diffuse across the spine cytoplasm to reach ATPases and other enzymes embedded in the spine head membrane. When these consumers are located farther from the mitochondrial source—such as on the distal membrane of a large mushroom spine head—the diffusion delay can become significant, potentially introducing local ATP gradients on short timescales. This is particularly relevant given the narrow diameter of the spine neck, which can restrict diffusion and prolong equilibration times [52, 53].

Furthermore, dendritic spines are dynamic structures: they undergo constant morphological remodeling and are sites of intense protein synthesis, turnover, and repair, especially following activity bursts or long-term potentiation (LTP) induction [9, 50]. These processes are all ATP-dependent. Local mitochondria, therefore, serve not only as energy suppliers for rapid electrophysiological recovery but also as metabolic hubs supporting longer-term biochemical and structural changes. Our findings support the concept that the coupling between mitochondrial positioning, calcium dynamic, and ATP production is a key determinant of synaptic efficiency. The tight regulation of ATP synthesis and delivery ensures that energy is not just available but spatially and temporally matched to local demands.

### 3.4 Metabolism in Soma versus Dendrites

Mitochondrial calcium uptake is compartmentalized in neurons, displaying distinct mechanisms in somatic versus dendritic regions. In the soma, blocking RyR-mediated CICR from the ER does not significantly impact mitochondrial Ca^2+^ transients. This suggests that, in cortical pyramidal neurons, somatic mitochondria primarily rely on direct calcium influx through voltage-gated calcium channels, particularly during action potentials, rather than on ER-derived calcium. High concentrations of IP3 receptors in the neuronal somata also reduce the dependence of mitochondrial function on the CICR mechanism.

These mitochondria appear tuned to integrate long-lasting or high-frequency electrical signals rather than localized synaptic activity. In contrast, we showed here that dendritic mitochondria exhibit a tighter functional coupling with ER calcium stores. This coupling is mediated through RyR-dependent CICR, which significantly enhances mitochondrial calcium uptake during coincident EPSPs and bAPs [54, 55]. Our findings support this view, showing that dendritic mitochondria positioned within 100 nm of RyR clusters at the spine base exhibit strong, rapid calcium uptake, leading to efficient ATP synthesis. This localized calcium amplification and spatial proximity to energy-consuming elements like ion pumps or synaptic signaling complexes allow dendritic mitochondria to meet the highly dynamic energy demands of synaptic activity. These differences point to a functional specialization: while somatic mitochondria may prioritize global energy homeostasis and are responsive to broader patterns of electrical activity, dendritic mitochondria are strategically positioned to supply energy for synapse-specific events. The localized calcium sensitivity of dendritic mitochondria supports rapid, spatially resolved ATP production, tightly coupled to synaptic transmission and plasticity.

### 3.5 How ATP Regulation Could Contribute to Synaptic Plasticity

Although we quantified here the time course of ATP dynamic during synaptic inputs, future directions should investigate how ATP production and delivery are modulated following long-term synaptic plasticity, such as LTP [14, 54]. Synaptic strengthening is known to increase metabolic demand due to elevated ion exchange, number of receptors, and local protein synthesis, suggesting that LTP necessitates enhanced mitochondrial ATP production to maintain homeostasis. Possibly LTP could be supported by structural or molecular remodeling of local mitochondria, including: (1) increasing the number or surface density of ATP synthase complexes on the mitochondrial face oriented toward the spine base; (2) relocating mitochondria to more proximal positions with respect to active spines; and (3) altering the organization of RyR-MCU microdomains to ensure reliable Ca^2+^-induced ATP production, although this last condition should be optimal. Such changes would enable mitochondria to dynamically meet the elevated energy demands associated with synaptic strengthening.

Moreover, there may be reciprocal signaling between spines and mitochondria—a form of “co-plasticity”—whereby increased synaptic activity not only drives LTP but also instructs nearby mitochondria to adjust their ATP output. This feedback could involve calcium-dependent signaling pathways, changes in mitochondrial membrane potential, or local translation of mitochondrial proteins. Notably, spine apparatus–bearing spines, which are more prevalent in mature, potentiated synapses, could serve as key loci for such metabolic adaptation due to their enhanced ability to amplify Ca^2+^ signals. Moreover, a new model of ATP production induced by Ca2+ would be helpfull to clarify the exact coupling ATP-Ca2+ coupling during synaptic plasticity[56]. Altogether, our findings suggest that synaptic activity is not merely a phenomenon of electrical and molecular signaling but also entails fine-tuned metabolic regulation. Future studies should integrate high-resolution imaging, metabolic sensors, and ultrastructural analysis to directly observe whether LTP/LTD induces changes in mitochondrial ATP production machinery and increase delivery in the spine head, and how this spatially correlates with the remodeled synaptic architecture.

## 4 Parameter values

## 6 Materials and Methods

### Ethics Statement

Animal handling was performed in accordance with the guidelines published by the Institutional Animal Care and Use Committee of the Weizmann Institute and with the Israeli National guidelines on animal care (IACUC), Israel (approval number: 00650120-3 from 20 January 2020 for 3 years).

### Culture Preparation

Cultures were prepared as detailed elsewhere [59, 60]. Briefly, E17 mouse embryos were removed from pregnant decapitated mothers’ wombs under sterile conditions, decapitated, and their brains removed. The hippocampi were dissected free and placed in a chilled (4 C), oxygenated Leibovitz L15 medium (Gibco, ThermoFisher Scientific, Inc., Waltham, MA, USA) enriched with 0.6% glucose and gentamicin (20 *µ*g/mL, Sigma, St. Louis, MO, USA). Tissue was mechanically dissociated with a sterile Pasteur pipette and passed to the plating medium consisting of 5% heat-inactivated horse serum (HS), 5% fetal bovine serum, and B-27 (1 *µ*l/1 ml, Gibco) prepared in minimum essential medium (MEM) Earl salts (Biological Industries, Beit Haemek, Israel), enriched with 0.6% glucose, gentamicin (20 *µ*g/ml), and 2 mM GlutaMax (Gibco) prepared in MEM-Earl salts. About 10^5^ cells in 1 mL medium were plated in each well of a 24-well plate onto poly-L-lysine (Sigma)-coated 13 mm circular glass coverslips. Cells were left to grow in the incubator at 37 °C, 5% CO_2_, and 3 days later the medium was replaced by FUDR (5-fluoro-2’-deoxyuridine, 50 *µ*g/ml, Sigma-Aldrich)-containing medium to block proliferating glia. The medium was replaced 4 days later by 10% horse serum in MEM after the transfection.

### Transfection

The transfection procedure was based and adopted from standard protocols [61, 62] and conducted at 8 days *in vitro* (DIV). Neurons were co-transfected with a genetically encoded FRET-based ATP biosensor, ATeam (pcDNA-AT1.03, dissociation constants for ATP ranging from 7.4 *µ*M to 3.3 mM), a gift from Prof. Hiromi Imamura, Kyoto University, Kyoto, Japan), which is comprised of a cyan fluorescent protein (CFP; mseCFP), an FoF1-ATP synthase *E* subunit and yellow fluorescent protein (YFP; cp173-mVenus) [63] and SP-short, subcloned into mCherry [64]. Lipofectamine 2000 (Thermo Fisher Scientific) mix was prepared at 1.2 *µ*l/well with 50 *µ*l/well OptiMEM (Gibco) and incubated for 5 min at room temperature. This was mixed with 1 *µ*g/well total DNA in 50 *µ*l/well OptiMEM and incubated for 15 min at room temperature (RT). The mix was then added to the culture wells and allowed to incubate for 3 h before a change of medium. Cultures were used for imaging at 11–24 DIV. Cotransfected cells displayed no apparent differences in spontaneous calcium activity, morphology, spine density, and survival. The distribution and pattern of the expression of the SP plasmid were similar to those of the endogenous SP [65]. The transfection protocol followed established methods [60].

### Ethics statement

Animal handling was performed in accordance with the guidelines published by the Institutional Animal Care and Use Committee of the Weizmann Institute and with the Israeli National guidelines on animal care (IACUC), Israel (approval number: 00650120-3 from 20 January 2020).

### Immunohistochemistry

#### Immunostaining

Cover glasses bearing transfected primary hippocampal cells were briefly washed with a standard extracellular solution and fixed with 4% paraformaldehyde in 0.1 M phosphate-buffered saline (PBS, pH 7.4) for 20 min. They were then thoroughly washed with PBS. Cultures were incubated for 1.5 h in 20% normal horse serum (NHS) diluted in PBS containing 0.2% Triton X-100. This was followed by overnight incubation at 4°C with primary antibodies: mouse monoclonal anti-ATPB (Abcam plc, Cambridge, UK, 1:500, gift from Prof. Atan Gross, Weizmann Institute of Science, Rehovot, Israel), and either rabbit anti-RyR (Abcam, ab2868, 1:250) or rabbit anti-MCU (Cell Signaling Technology, D2Z3B, #14997, 1:500), depending on the experiment.

After primary antibody incubation, cultures were incubated for 1.5 h with secondary antibodies: Cy2-conjugated anti-mouse (1:250) and Donkey Anti-Rabbit IgG H&L (DyLight^@^ 650, 1:250). Coverslips were then rinsed with PBS, mounted onto glass slides, and visualized using a Zeiss LSM 880 upright confocal microscope (Oberkochen, Germany) equipped for simultaneous imaging of four fluorophores, using a Plan-Neofluar 40*×* /1.30 oil immersion objective and an anti-fade mounting medium.

### Chemicals Application

Cell cultures were incubated with Fluo-2 AM (2 *µ*M, ION Biosciences, Texas, USA) for 1 hour at room temperature, or X-Rhod-1 AM dye (1.5 µM, Thermo Fisher Scientific) for 30 minutes, to image fluctuations in cytosolic calcium concentration [Ca^2+^]_c_ in standard recording medium containing (in mM): NaCl 129, KCl 4, MgCl_2_ 1, CaCl_2_ 2, glucose 10, HEPES 10, pH adjusted to with NaOH and osmolality to 320 mOsm with sucrose.

### Imaging

We used the lambda mode of a Zeiss LSM880 laser scanning confocal microscope equipped with a 32-channel GaAsP detector array, which allows fluorescence emission spectrum splitting into spectral intervals recorded in individual detection channels. CFP/YFP-paired FRET (MBS T80/R20) and Fluo-2 were excited with the 458 nm argon laser, SP-mCherry or X-Rhod-1 were excited with the 543 nm Helium-Neon laser, and the fluorescence detection range was set between 460 nm and 701 nm with a spectral step of 8.9 nm per channel. Lambda stacks for each fluorophore were pre-acquired using a 40x 1–numerical aperture (NA) water immersion objective Plan-Apochromat.

### Data Analysis

Linear unmixing software tool (ZEN Microscopy Software, Zeiss, Germany) was used for spectral unmixing based on spectral profiles from pre-acquired Lambda stacks. The signal from SP-mCherry was subtracted from the analysis by linear unmixing and was not used in subsequent analysis. Fluorescence intensity was calculated using ZEN (Zeiss, Germany), ImageJ (NIH, Bethesda, MD, USA), MATLAB (MathWorks Inc., Natick, MA, USA), and Origin Pro 2021 (Electronic Arts, Inc., San Mateo, CA, USA) software.

### Stochastic model of calcium dynamic and numerical simulations

In our simulation, we investigate two geometries of the spine, the mushroom and stubby ones (spherical head connected to the dendrite by a cylindrical neck). The details of the geometries are summarized in Table 1. The bottom of the SA is always aligned with the bottom of the spine neck. A spherical mitochondrion with radius *R*_*mito*_ = 0.4 *µ*m is located below the bottom of SA with the gap *d*_*gap*_. The radius and length of the dendritic shaft are 0.585 *µ*m and 5 *µ*m, respectively.

#### Stochastic Modeling and simulations of Calcium Diffusion and Reactions in Dendritic Spines

To simulate the nanometer-scale spatiotemporal dynamic of Ca^2+^ ions in dendritic spines, we employed a Brownian dynamic framework based on the Euler–Maruyama scheme [35]. Ca^2+^ ions are modeled as independent Brownian particles following the overdamped Langevin equation:

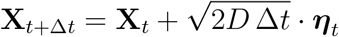

where *D* is the effective diffusion coefficient of Ca^2+^ (typically 600 *µ*m^2^*/*s), Δ*t* is the time step, and ***η****t ∼𝒩* (0, **I**) is a standard Gaussian noise vector. The time step is set to Δ*t* = 1 *µ*s, which is much shorter than the binding and releasing behavior of the calcium buffers, RyR and MCU. To simulate Ca^2+^ diffusion in the 3-D space within the spine and dendrite, the surface of the spine, SA, and mitochondrion are treated as a reflecting boundary for diffusion. The two ends of the dendritic shaft are set as an absorbing boundary.

**Geometry**. The simulation domain consists of:

- A spherical spine head (*R ≈* 0.4 *µ*m),
- A cylindrical spine neck (*R*_*n*_ *≈* 0.15 *µ*m, *L*_*n*_ *≈* 0.5 *µ*m),
- A dendritic shaft (radius *≈* 0.585 *µ*m, length 5 *µ*m).

#### SERCA Pumps

SERCA pumps are modeled as small absorbing disks of radius 10 nm located on the spine apparatus (SA). A SERCA binds a single Ca^2+^ ion and enters a refractory state for 100 ms, during which it is non-interacting. SERCA pumps are modeled as 50 small targets uniformly distributed in the upper surface of the spine constituting a radius of 10nm each. Note that we do not model here the SERCA pumps which have little impact on the initiation dynamic of CICR ([35]).

The interaction with RyR Receptors, MCU and CaM buffer is described below.

**Initial Conditions for Releasing Ca**^2+^. The three different release locations are modelled:

1. **Synaptic input**: 1000 Ca^2+^ released instantaneously at the top of the spine head;
2. **Upper bAP**: 1000 Ca^2+^ ions released at the upper point of the dendritic shaft with the 8 *µ*m distance to the spine center.
3. **Lower bAP**: 1000 Ca^2+^ ions released right below the mitochondrion at the lower point of the dendritic shaft.

In the simulation, we do not consider any interactions between the calcium ions.

#### Numerical Implementation

Simulations were implemented in MATLAB, tracking the full stochastic trajectories of up to 2000 particles. Absorbing interactions (e.g., with RyR, SERCA, CaM, MCU) are resolved by checking for proximity within a specified catchment radius. Time-resolved binding curves and histograms are generated over 10–20 realizations for statistical robustness [35].

#### Simulations of RyR activitation

Ryanodine Receptors (RyRs) are calcium-gated channels located on the membrane of the spine apparatus. Their activation follows a CICR mechanism: the stochastic biding of Ca2+ to RyRs follow the methodology introduced in [35]. RyRs are located in line as previously and not overlap with the spherical mitochondrion. We positioned 36 RyRs (e.g., 3–4 RyR rings) in four horizontal rings within a 250nm range at the base of the spine. 37.5 nm for the mushroom and 87.5 nm for the stubby.

- **Activation Rule:** Each RyR channel opens only upon the arrival of two calcium ions within a time window of 7.5 ms. When the binding radius of RyR is 10 nm. The first ion primes the receptor, and the second triggers its opening.
- **Calcium Release:** Upon activation, the RyR channel releases a fixed number of calcium ions from the ER into the cytosol. In our simulations, each RyR releases 10 *−* 2*N* Ca^2+^ ions along with the two bound ions, after a delay of 0.25 ms after the second Ca2+ binding, and *N* is the number of times that the RyR has released. The released ions will be placed at random locations where 20 nm away from the RyR center.
- **Refractory Period:** Refractory Period: After release, the RyR channel enters a refractory state lasting 4.5 ms, during which it is non-responsive to further calcium and modeled as a reflective surface for free Ca^2+^.
- **Spatial Organization:** In the simulation geometry, RyRs are positioned at the base of the spine apparatus and oriented toward the mitochondrion. They are arranged in 4 concentric rings with 9 RyRs each, totaling 36 RyR channels. The distance between RyRs and mitochondrial calcium uniporters (MCUs) can vary starting with a minimal distance of 20 nm.

#### Modeling and Simulations of MCU Activation by Calcium Ions

Mitochondrial Calcium Uniporters (MCUs) are calcium-selective channels located on the outer membrane of mitochondria, responsible for calcium-dependent ATP production. In our stochastic simulations, we model MCU activation based on calcium binding kinetics and nanometer-scale spatial constraints:

- **MCU Geometry and Placement:** The mitochondrion is modeled as a sphere with a radius 0.4 *µm*, positioned adjacent to the base of the dendritic spine apparatus. MCU channels are treated as small absorbing disks (binding sites) located on the mitochondrial surface. We explore two spatial distributions:
- **Polarized distribution:** The locations of 50 MCU channels are generated randomly on the upper hemisphere of the mitochondria (facing the spine base and ER).
- MCU channels are distributed 1-Upper half only and 2-uniformly over the entire mitochondrial surface.
- **Binding Rule:** Each MCU channel is modeled as requiring **three calcium ions to bind within a time window of 20ms** in order to transition into the activated state. The first and second Ca^2+^ ions that arrive are stored temporarily. If the third ion does not arrive within 20ms of the first, all bound ions are released, and the channel resets.After the release of ions, the binding site will experience a refractory period of 1 ms.
- **MCU Activation State:** Upon successful binding of 3 calcium ions, the MCU is marked as active and remains permanently in this state for the remainder of the simulation.
- **MCU Catchment Radius:** Each MCU channel is modeled with a catchment (binding) radius of 10nm.
- **MCU deactivation:**After 60 ms of activation of a MCU channel, the three bound Ca^2+^ will be released and the channel will be closed with a refractory period of 1 ms before it can be reactivated.
- **Output Metrics:** At each time point, we record the total number of MCUs that have successfully transitioned to the activated state in both synaptic and bAP scenarios.

#### Stochastic modeling and simulation of calcium binding to calmodulin (CaM)

We simulate the activity of calmodulin, the major endogenous calcium buffers binding to calcium in the spine and the dendrite. Calmodulin (CaM) is modeled as a molecule composed of two lobes, the N-terminal (N-lobe) and the C-terminal (C-lobe), each of which can bind up to two Ca^2+^ ions. The lobes exhibit different calcium binding kinetics: the N-lobe binds and unbinds Ca^2+^ more rapidly and with lower affinity, while the C-lobe binds more slowly but has higher affinity and a longer residence time.

In our stochastic simulations, Ca^2+^ ions are represented as Brownian particles. A Ca^2+^ ion can bind to a lobe when it diffuses within a spherical *catchment radius* of *r* = 10 nm from the center of the lobe. Each CaM lobe is modeled as having two independent binding sites. When a Ca^2+^ ion enters the catchment region and the lobe has available binding sites, the ion is bound with probability 1 (i.e., immediate capture). Once bound, each Ca^2+^ ion remains associated with its respective lobe for a random dwell time drawn from an exponential distribution governed by the lobe-specific unbinding rate *k*_off_:

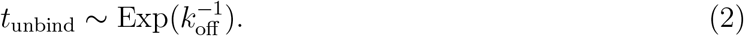

The parameters used are:

- **N-lobe:** *k*_off,N_ = 100 s^*−*1^ (mean unbinding time *∼*10 ms)
- **C-lobe:** *k*_off,C_ = 1 s^*−*1^ (mean unbinding time *∼*1 s)
- **Binding radius:** *r* = 10 nm
- **Number of CaM molecules:**200 CaMs are placed in the mushroom spine (volume 0.30*µm*^3^) and 225 CaMs in the stubby spine (volume 0.33*µm*^3^). We placed 1,500 CaMs in a region measuring 1.6 *µ*m in length along the dendrite, centered on the mitochondrion. The volume of the region is 1.45*µm*^3^. To save computation, we does not place the buffers in the whole dendrite. The locations of the buffers are generated randomly in the corresponding regions. We generate 1700 in a volune of *V* =_*d*_= 1.45*µm*^3^ (dentrite) and 120 in 0.3*µm*^3^ (spine), thus accounting for the factor 3 difference in concentration between the dendritic spine and dendrite [37].
- **release of ions:** After *t*_unbind_ of binding, the bound ions will be released and placed at a random location 20 nm away from the center of the lobe.

Each simulation trial records the number of Ca^2+^ ions bound to each lobe type over time. Final results are averaged over multiple stochastic realizations (typically 10) to account for inherent molecular variability. In our simulations, each calmodulin (CaM) molecule is modeled as a linear two-lobed structure, comprising an N-terminal lobe (N-lobe) and a C-terminal lobe (C-lobe). Each lobe is represented by a spherical binding center with a catchment radius of 10 nm for calcium (Ca^2+^) binding. The two lobes are arranged along a common axis (x-axis), separated by a fixed surface-to-surface distance of 4 nm. This linear geometry approximates the extended conformation of calmodulin observed in its calcium-bound state. We took a distance between the N-lobe and C-lobe of 4 nm. This separation reflects structural crystallography data [66, 67, 68]. Each lobe is independent and detects and binds Ca^2+^ ions stochastically, based on local proximity and lobe-specific unbinding rates. Note that we chose 1700 Calmodulin to account for possible endogeneous already bound calcium.

#### Modeling and simulations of ATP interacting with exchangers

We computed the probability and the mean time for ATP molecules generated at some position in the dendrite to arrive to exchanger sites located on the spine head surface. For this goal, we model ATP motion as diffusing and thus solve the associated equation (section S7) and also use Comsol multi-physics when adding the mitochondria ball geometry.

##### Time scale of ATP to exchangers located in a dendritic spine

We simulate ATP in a dendritic spine (spine head, neck, and dendrites parameters are given in Table S1). Brownian ATP molecule follows the stochastic equation 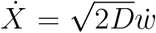, where *w* is the Wiener white noise. The ends of the dendrite are modeled as absorbing boundaries, thus molecules arriving at the base of the spine do not re-appear in the simulation. ATP can absorbed by the two sides of the dendrite (ATP molecule moving in the dendrite will not reenter the spine neck in a reasonable time compared to ATP entering directly).

Exchangers are distributed on the spine head upper hemisphere. Exchangers are modeled as absorbing circular disks with a catchment radius *A* = 10*nm*. After the first ATP is bound, the target exchanger is no longer available for binding and thus acts as a reflecting boundary. This simulation scheme is implemented in Fig. 5.

### 6.1 Computing the time for ATP molecules to reach exchangers located on the spine head surface

The time for ATP molecules to reach an exchanger located on the spine head surface is computed from formula [43, 69]

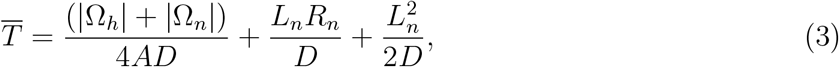

where |Ω_*h*_| = 4*πR*^3^*/*3 and 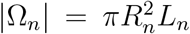 are the neck and head volumes. The term 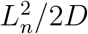 accounts for the diffusion time across the neck, while *L*_*n*_*R*_*n*_*/D* models the mean time for an ATP molecule to cross from the neck to the head. There are *n*_*r*_ exchangers.

Another quantification to estimate exchanger activation is provided by the formula for a single ATP bind to bind one of the *n*_*r*_ exchanger:

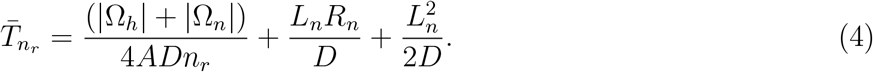

For a total of *N* diffusing ATP molecules that need to bind all *n*_*r*_ receptors (where *N ≥ n*_*r*_), the mean time 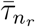 is computed as the last passage time of *N − k* available ATP molecules to the remaining *n*_*r*_ *− k* receptors, Fig. 5. The mean time [43, 70] is given by

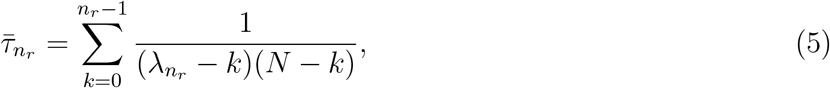

where 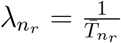

### 6.2 Splitting probability formula for ATP between spine and dendrite

Computing the splitting probability for a Brownian particle is equivalent to solve the Laplace equation in the spine-dendrite with appropriate boundary conditions [43]. We present now the splitting probability, which consists in computing the fraction ATP molecules entering the head versus the ones lost in the dendrite. A thorough computation [69] leads to the formula

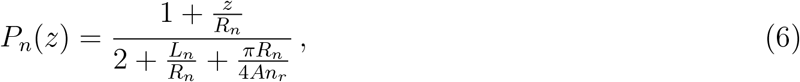

where *z* is the distance of the initial ATP initial release site from the bottom of the neck, *R*_*n*_ and *L*_*n*_ are the radius and length of the neck respectively, while there are *n*_*r*_ exchangers of radius *A*. The probability *P*_*n*_(*z*) accounts for the fraction of ATP that will reach exchangers in the spine head before being lost in the two sides of the dendrite. Typical spine neck length is *L*_*n*_ = 500nm and the head radius is *R*_*n*_ = 150nm as used in the simulations.

The probability *P*_Exchangers_(0) that an ATP molecule released at the base of the spine enters the head (in the absence of cytosol consumers) and binds one of the *n*_*r*_ membrane-bound exchangers before escaping into the dendrite [69]:

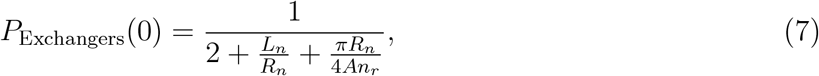

where *L*_*n*_ and *R*_*n*_ are the length and radius of the spine neck, *A* is the capture radius of an individual exchanger (typically 10 nm), and *n*_*r*_ is the number of exchangers. We use here the representative values (*L*_*n*_ = 0.5 *µ*m, *R*_*n*_ = 0.15 *µ*m, *A* = 0.01 *µ*m, and *n*_*r*_ = 50) leading to *P*Exchangers(0) = 18%.

### Quantifying RyR, ATP synthase and MCU in mitochondria and ER in dendrites

To quantify the level of expression and organization of ATP concentration, ATP synthase and MCU expression in mitochondria relative to RyR in the ER, we develop the following segmentation procedure: we collected data from ATP-synthase, calcium concentrations in dendritic spine head and in the corresponding dendritic spines for 24 dendritic spines with varying lengths from 0.17*µm* to 1.15*µm*. RyR and ATP-synthase molecules are collected from different regions of interests (ROIs) *R*_1_, *R*_2_, *R*_3_, *R*_4_ for 71 dendritic mushroom and 13 dendritic stubby spines across 3 experiments displaying different lengths and head diameters in the range 0.24 *µm* to 2.00 *µm* and from 0.19 *µm* to 1.41 *µm* respectively for mushroom spines, and from 0.18 *µm* to 0.55 *µm* and from 0.20 *µm* to 0.88 *µm* respectively for stubby spines.

For mushroom and stubby spines, the regions *R*_*i*_ associated to ATP-synthase provide a partition of the adjacent mitochondria located at the basis of dendritic spines divided into upper and lower region, aligned and a bit a side.

For mushroom and stubby spines, the regions *R*_*i*_ associated to RyR are located in the spine neck, and in the top and bottom parts of the dendritic spine, adjacent to spine head. As a control region for RyR and ATP-synthase, we also use a ROI *R*_5_ selected at a mitochondrial region far from adjacent spines (Fig. 3B)

#### Model Evaluation Metrics

The spatial distribution analysis of RyR and ATP-synthase expression in mitochondrial regions of interest (ROIs) near mushroom and stubby spines (fig. **B**) to estimate if distribution were different at different distances from the spine head. To quantify the spatial distribution, we define ROI *R*_*i*_, *∀i ∈ {* 1, 2, 3, 4, 5*}* and displayed in red for ATP-synthase, in blue for RyR. To measure if the distribution decrease with the distance from spine head was significant from on region to another, a paired one-tailed Wilcoxon signed-rank test is applied between each specific region and the relative reference one regarding the marker (ATP-synthase or RyR represented by bi-directional arrows on image). The significant differences were noted for various regions, with the respective p-values indicating different levels of significance compared to the reference region: * : *p <* 0.05, ** : *p <* 0.01, *** : *p <* 0.001, **** : *p <* 0.0001. For each distribution, the relative importance of the distribution of the specific region compared to the reference one is provided as a percentage in magenta for ATP-synthase and green for RyR.

#### Regression procedures

To quantify the level of ATP and ATP-synthase expression vs spine length *L*_*spine*_ and spine head diameter *Diam*_*Head*_ or RyR vs *Diam*_*Head*_, we use a polynomial regression of degree 3 using the function

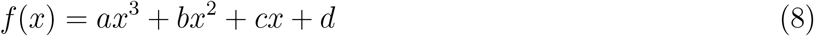

For *Ca*^2+^ level in dendrite vs the spine lenghth, we fit a sigmoid

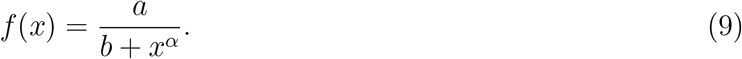

For RyR and ATP-synthase against spine length and spine head diameter, we perform a simple linear regression. Parameters 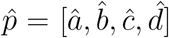 for polynomial, 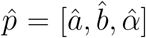 for sigmoid are obtained from a least-square optimization

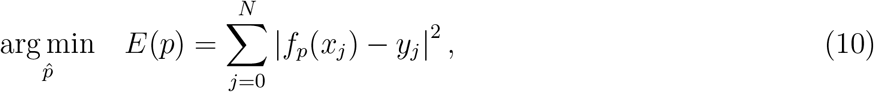

where *N* is the number of data samples, (*x*_*j*_, *y*_*j*_) are the data samples, and *f*_*p*_ is the function of parameter *p* used for the regression and *E* is the residual error of the least-square problem. We use for polynomial fitting .**polyfit()** from the numpy Python library, the sigmoid regression is performed using .**curvefit()** from the scipy.optimize Python library.

#### Model Fitting

To assess the statistical relevance of the regressions, we use three indicators: *R*^2^, F-statistic, and *p*_*value*_, derived from the ANOVA table. These indicators are calculated as follows:

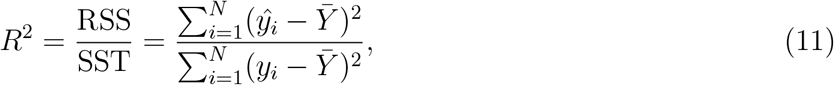

where SST stands for “Total sum of squares” corresponding to the variations of the data relative to their mean, and RSS stands for “Regression Sum of Squares” corresponding to the total variations of the data explained by the regression model. The *R*^2^ is directly related to the Pearson correlation coefficient as 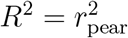. We use the F-statistic to compare the variances of the data with the one of the model:

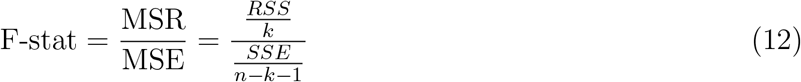

where *n* is the number of data samples and *k* is the number of degrees of freedom of the regression model. SSE is the Sum of Squared Errors, i.e., 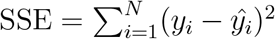. A high fit and F-stat (rule of thumb *>* 4) is associated to a model that accounts for a substantial proportion of the variance of the data. The *p*_*value*_ is computed from the F-stat and the degrees of freedom as:

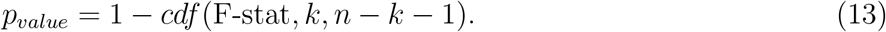

A regression model accounting well for the data correctly has a high *R*^2^, a high F-stat, and a low *p*_*value*_. The regression results are the following:

1. ATP vs *L*_*neck*_ we obtain for the polynomial (*a, b, c, d*) = (0.33, *−*1.97, 1.92, 0.06), with (*R*^2^ = 0.55, F-stat = 5.72, *p*_*val*_ = 3.4 *×* 10^*−*3^).
2. Ca2+ concentration vs *L*_*neck*_, the coefficients are (*a, b, α*) = (0.36, 0.37, 2.68), with significance (*R*^2^ = 0.55, F-stat = 8.22, *p*_*val*_ = 9.27 *×* 10^*−*4^)
3. RyR distribution vs spine length has been fitted by a sigmoid function equation 9 with parameters 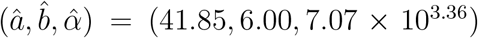. The fit is significant since it provides (*R*^2^, *F*_*stat*_, *p*) = (0.30, 12.54, 7.07 *×* 10^*−*7^).
4. RyR vs *Diam*_*Head*_ is fitted by a degree 3 polynomial function with equation 9 with parameters 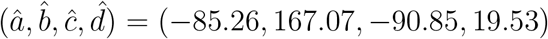. The fit is significant since it provides (*R*^2^, *F*_*stat*_, *p*) = (0.12, 2.92, 2.61 *×* 10^*−*2^).
5. ATP-synthase vs *L*_*neck*_ fitted by a degree 3 polynomial function with equation 9 with parameters 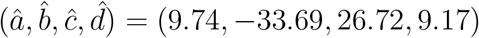 with (*R*^2^, *F*_*stat*_, *p*) = (0.51, 28.88, 2.22 *×* 10^*−*16^).
6. ATP-synthase vs spine head diameter has been fitted by a degree 3 polynomial function with equation 9 with parameters 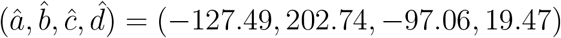 with (*R*^2^, *F*_*stat*_, *p*) = (0.31, 11.91, 4.69 *×* 10^*−*8^)

The distribution of ATP-synthase and RyR decrease both with the distance from spine head for mushroom or stubby spines. This is significant as shown by the Wilcoxon test as summarized in table 3 below:

**Table 3.**
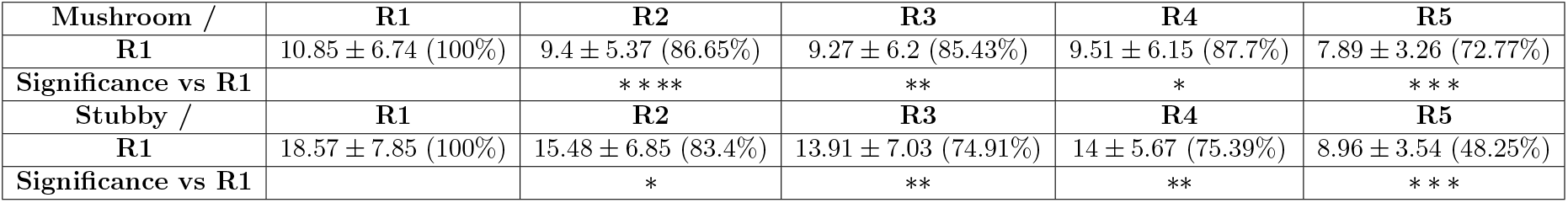
Distributions of ATP-synthase for Mushroom vs Stubby.

**Table 4.**
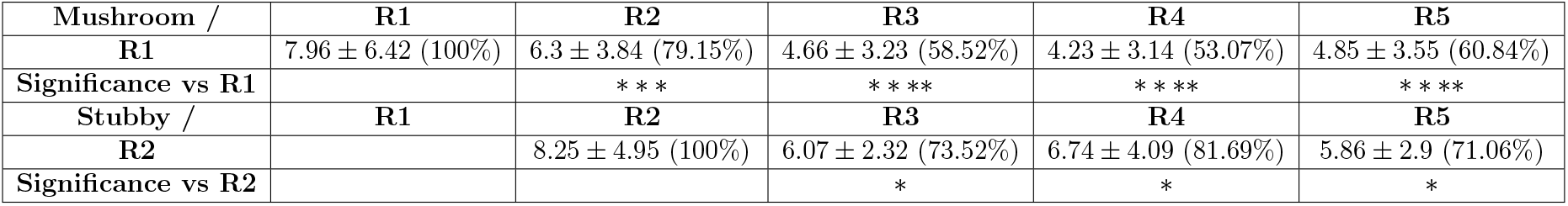
Distributions of RyR for Mushroom vs Stubby.

